# The neural mechanisms of fast versus slow decision-making

**DOI:** 10.1101/2024.08.22.608577

**Authors:** Mostafa Nashaat, Hatem Oraby, Flutra Krasniqi, Sek Teng Goh-Sauerbier, Marion Bosc, Sandra Koerner, Sedef Karayel, Adam Kepecs, Matthew E. Larkum

## Abstract

Is more haste less speed? Decision time variability has been attributed to speed/accuracy trade-offs(1), internal mental architecture(2), and noisy evidence accumulation(3–5). However, exploring these possibilities is difficult in rodents that consistently behave impulsively. Here, we demonstrate a novel floating-platform system that allows head-fixed mice to voluntarily vary decision times, akin to observed human behavior, in combination with sensitive neuroimaging approaches. We track the activity flow from medial to lateral frontal cortex (MFC to LFC) and record sequences of single-neuron activity. Choice-selective neurons displayed divergent temporal codes between MFC and LFC, with remarkable MFC susceptibility to optogenetic inhibition. These results suggest that LFC acts as an integrative motor threshold, while MFC plays a broader cognitive role in strategy and choice-selection.

## Results

During natural decision-making paradigms in humans, there are many variables at play such as task history (priors), task difficulty (evidence), stakes (cost/reward) and attention(*3*, *5*, *6*). To disentangle the causal neural mechanisms underlying these factors, controlled experimental conditions are necessary and best studied in head-fixed mice. In this study, we developed a head-fixed mouse preparation combined with a floating nose-poke platform technology (“AirTrack”(*7*)), enabling the use of advanced neural acquisition and perturbation techniques, while simultaneously facilitating more natural, quasi-freely-moving behavior(*8*, *9*). Head-fixed mice voluntarily navigated a nose-poke behavioral environment on a floating platform (Fig. 1A, B; fig. S1A), to perform a two-alternative forced-choice visual discrimination task (Video S1). To support self-paced evidence accumulation, we utilized the classical Random Dots Kinematogram (RDK) with free sampling adapted for rodents(*10–12*)(fig. S1B; “free sampling time”, see methods). Mice were rewarded by moving in the direction that best matches the movement of the dots. By varying the dots’ coherence, we could finely control stimulus salience and trial difficulty.

**Fig. 1.**
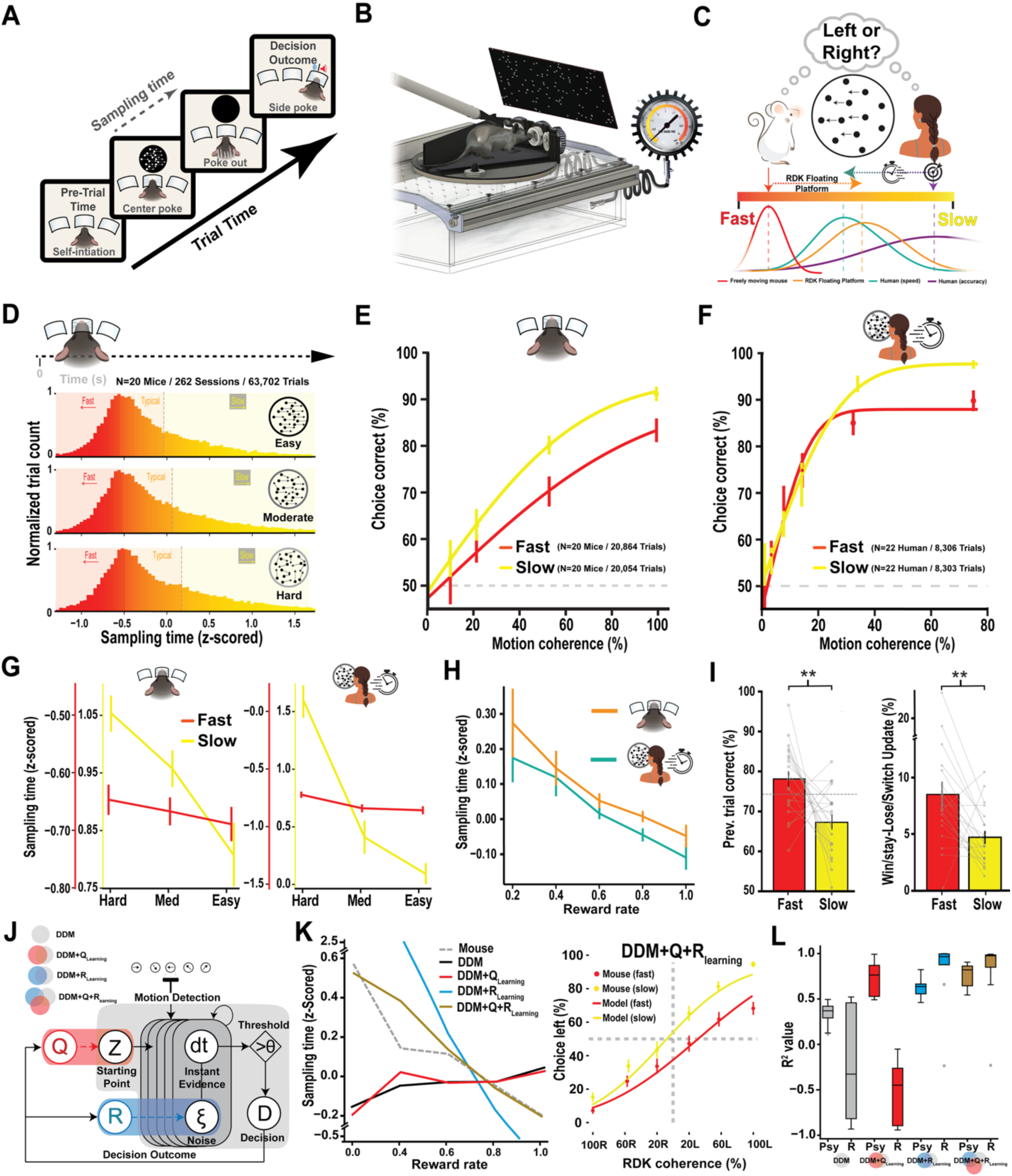
A novel behavioral paradigm demonstrates similar adaptive decision-making strategies in mice and humans. (**A**) Schematic of the Random Dots Kinematogram (RDK) task for head-fixed mice using a nose-poke floating platform. (**B**) 3D illustration of the floating-platform apparatus with three nose-pokes designed for voluntary decision-making tasks in head-fixed mice. (**C**) Comparison of sampling time distributions in mice and human highlighting flexible decision-making strategies across different contexts, including freely moving mice performing a visual detection task (red), head-fixed mice on the floating platform performing the RDK task (orange), and humans performing the RDK task under accuracy condition (violet) or speed condition (turquoise) instructions. These distributions show a continuum from impulsive (“fast”) to deliberate (“slow”) decision-making. (**D**) Histograms of sampling times categorized into fast (red), typical (orange), and slow (yellow) trials across stimulus difficulties (easy, moderate, hard). (**E**) Performance accuracy (% correct choices) in head-fixed mice as a function of task difficulty (motion coherence %) for fast (red) versus slow (yellow) trials. (**F**) Human performance accuracy during speed-instructed RDK trials, formatted as in panel E. (**G**) Mean sampling time (z-scored) of correct trials across stimulus difficulties during fast (red) and slow (yellow) trials for both head-fixed mice and human subjects (speed condition). (**H**) Mean sampling time (z-scored) as a function of recent reward history (reward rate calculated from the preceding 5 trials) in both head-fixed mice and humans (speed paradigm), illustrating behavioral sensitivity to reward feedback. (**I**) Left: Probability of adopting fast versus slow strategies based on previous trial outcomes (correct/incorrect) in mice. Right: Win-stay/Lose-switch behavioral update probability during fast versus slow decisions in mice. (**J**) Schematic representation of reinforcement learning (RL)-enhanced drift diffusion models (DDM) tested to capture mouse behavior. (**K**) Left: Reward rate plotted against z-scored sampling time from an example mouse (avgat_1 = 2501 trials; dashed gray line) compared to predictions from different RL-enhanced DDM variants: basic DDM (black), Q-learning combined DDM (DDM+Q, red), R-learning combined DDM (DDM+R, blue), and Q-learning & R-learning combined DDM (DDM+Q+R, gold). Right: Psychometric curve of the same example mouse for fast (red dots) and slow (yellow dots) trials, along with fits from the Q&R-DDM model (lines). (**L**) Pearson correlation (R²) values for model fits (DDM, DDM+Q, DDM+R, DDM+Q+R) and mice behavioral measures: psychometric performance for fast and slow conditions (“Psy”), and sampling time/reward rate sensitivity (“R”).

Unlike humans, freely moving mice performing simple detection tasks often display impulsive behavior under commonly used approaches even when sampling time is unrestricted (Fig. 1C, red histogram; fig. S1C)(*13*, *14*). To address mice impulsivity, we adapted the RDK task to a floating platform, encouraging more voluntary control over sampling time and thus revealing richer decision-making patterns (Video S2). We have also conducted a comparable experiments in human subjects that involved speed-accuracy trade-off adjustment, adding a competitive time constraint to mirror the task design used in mice(*15–17*) (fig. S2A, see methods). In humans, this manipulation produced a clear shift from slow and accurate to fast and impulsive behavior (Fig. 1C, Speed = turquoise, n=20; Accuracy = violet, n=22; fig. S1C; fig. S2B). The combined reciprocal adjustments allowed us to align the human and mouse tasks, making their behavioral outputs directly comparable (fig. S2C).

Having established task comparability, we next asked what could be learned from the way sampling times vary across trials and conditions (Video S3). We began by dividing the sampling time distribution according to trial difficulty (‘easy’, ‘medium’ & ‘difficult’; see methods). In mice, examining the trial-time distributions across all trials grouped by difficulty, revealed that task difficulty had surprisingly little influence on sampling time (Fig. 1D). We then divided sampling times within each difficulty into thirds, labeling trials in each segment as fast (’impulsive’, red), typical (’typical’, orange), or slow (’deliberate’, yellow) (Fig. 1D; n=20, session=262, trials=63,702 trials; example mouse, fig. S2D&E). Performance scaled with both task difficulty and sampling duration, increasing with both easy and slow trials (All mice, Fig. 1E; example mouse, fig. S2F). Next, we compared psychometric curves across fast and slow trials (Fig. 1E). While both followed the expected dependence on stimulus difficulty, slow decisions yielded significantly higher accuracy than fast ones (All mice, Fig. 1E; example mouse, fig. S2F). The same pattern was observed in humans performing the task in the speed context (Human speed: Fig. 1F, fig S2G&H; human accuracy: fig S2I-K), with fast trials showing significantly higher lapse rate than slow ones (fig S2L).

Because fast and slow trials differed in performance, particularly on easier tasks, we restricted our analysis to correct trials to examine whether sampling time reflected an underlying decision strategy (Fig. 1G). In doing so, we observed a shift in policy for both mice and humans (speed paradigm): subjects voluntarily increased sampling time for difficult trials only during slow (deliberate) trials, but not during fast (impulsive) trials (mice, Fig. 1G, left: fast, slope= 0.02, θ=1.08°; slow, slope=0.14, θ=7.83°; human speed, Fig. 1G, right: fast, slope= 0.06, θ=3.70°; slow, slope=0.74, θ=36.52°; human accuracy, fig S3A). Voluntarily adjusting the time to consider the evidence during slow trials but not fast trials suggests that factors other than difficulty can sometimes influence behavioral strategy(*18–20*). We hypothesized that subjects update their decision-making strategy to maximize an objective function, e.g. perceived reward outcome(*21–23*). Here, we found that the average reward rate, i.e., performance over the last 5 trials, significantly influenced sampling time in both mouse and human tasks (Fig. 1H, fig. S3B–D). Thus, there was a tendency in mice to slow down if the previous trial was incorrect (Fig. 1I, left; fig. S3E, F)(*24*). Furthermore, during fast decisions, mice were significantly biased to switch sides if the previous trial was incorrect and a corresponding bias to stay if the previous trial was correct (Fig. 1I, right, fig. S3G-I). This indicates that fast and slow behavioral strategies arise from distinctively different behavioral outcomes in preceding trials. We found that previous reward history had the highest explanatory power (34.36%±9.94SEM), followed by evidence (27.15%±6.87SEM), trial history update (21.02%±5.82SEM) and motor bias (19.32%±5.46SEM; fig. S3J; see methods).

We next explored what underlying computations might account for the variation in sampling time and decision strategy. Drift-diffusion models (DDMs) are commonly used to explain sampling time, but on their own, we found that they fell short of capturing key aspects of strategic variability, particularly those linked to reward history(*4*, *25*) (Fig. 1J; fig. S4A). This led us to ask whether reinforcement learning components, which incorporate trial history, might improve model predictions when combined with DDM(*25*, *26*) (Fig. 1J, K). We tested two reinforcement learning models: Q-learning, where reward sites are represented as states with associated action values, and R-learning, where the average reward rate modulates integration noise (fig. S4A&B; see methods). Only when both Q- and R-learning were combined with the classical DDM could we best capture the behavioral phenotypes observed in mice, including the effects of reward rate on sampling time and the fit of the psychometric function (Fig. 1J-L; “DDM+Q+R”).

### Spatiotemporal recruitment of frontal cortex mediates behavioral strategy

To investigate the cellular mechanisms of fast versus slow decisions, we exploited the floating platform approach in combination with widefield imaging and optogenetic perturbation of specific cortical regions. For widefield calcium imaging, we used a transgenic mouse line (Thy1 GP4.3) to express a genetically encoded calcium indicator (GCaMP6s) in cortical excitatory neurons (Fig. 2A; and fig. S5A; see methods)(*27*), while mice performed the RDK task using the voluntary-sampling paradigm (free sampling; Fig. 1A). Given the variable nature and timing of behavioral strategies (Fig. 1D), we first focused on typical behavioral trials (sampling times of 1.0-1.2s; fig. S5B; see methods) while using widefield imaging to characterize activity across the cortex (Fig. 2B&C; fig. S5C&D)(*28*). We observed that all areas around the cortex were active at some point during the sampling with a distinct spatio-temporal pattern originating from the medial axis of the dorsal neocortex at the start of sampling (Fig. 2B, left), and traversing radially until the animal made a decision (Fig. 2B and fig. S5E, decision made after last panel).

**Fig. 2.**
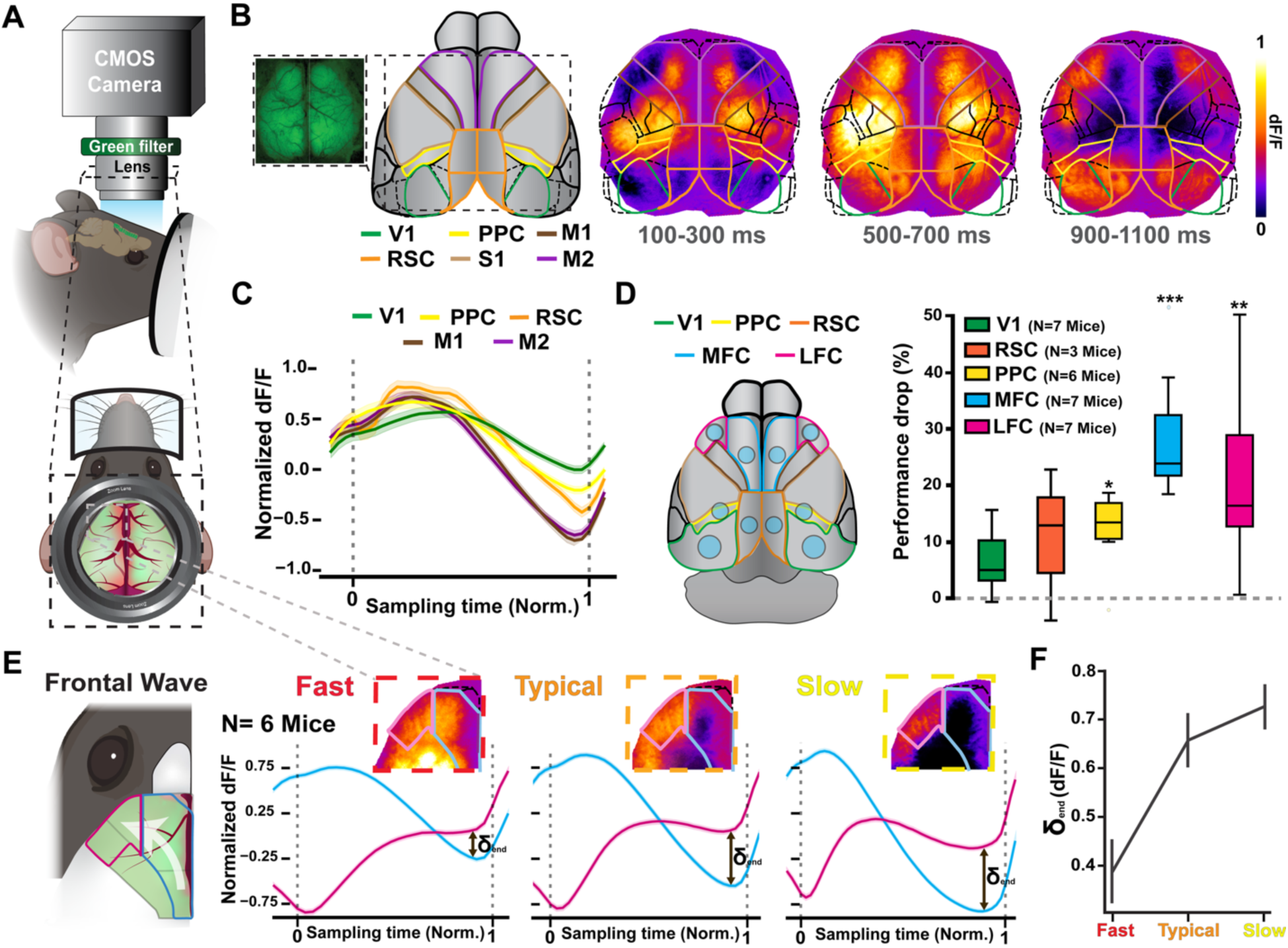
Spatiotemporal activation of frontal cortex regions causally relates to fast and slow decision-making strategies. (**A**) Schematic illustrating the widefield calcium imaging setup used in head-fixed mice. (**B**) Pixel-wise heatmaps depicting average calcium fluorescence (dF/F) in an example Thy1-GCaMP6s mouse across three different time windows during a typical sampling interval (1.0–1.2 s) in the RDK decision-making task. Cortical area boundaries are outlined for reference according to Allen brain atlas. (**C**) Mean normalized calcium responses across different cortical areas indicated in panel B (primary visual cortex, V1; posterior parietal cortex, PPC; retrosplenial cortex, RSC; primary motor cortex, M1; secondary motor cortex, M2), averaged across 6 sessions in 4 mice for trials between 1 to 1.2 seconds. Shaded regions represent SEM. (**D**) Behavioral impact of bilateral optogenetic silencing (using VGAT-ChR2 mice) targeting distinct cortical regions during a fixed, 1-s sampling RDK task. The cortical map illustrates targeted areas: V1 (visual), PPC (posterior parietal), RSC (retrosplenial), MFC (medial frontal cortex), and LFC (lateral frontal cortex). Boxplot quantifies the percentage drop in task performance following cortical silencing (**\***p < 0.05, **\*\***p < 0.01, \***\*\***p < 0.001). (**E**) Functional segmentation of frontal cortex activity into medial frontal cortex (MFC; blue) and lateral frontal cortex (LFC; magenta), based on distinct spatiotemporal activation patterns during the sampling period. Representative heatmaps illustrate average calcium activity patterns for fast (impulsive), typical, and slow (deliberate) decision-making strategies. Corresponding normalized calcium fluorescence signals for MFC and LFC (n=6 mice) are plotted below each heatmap. The double-headed arrow labeled “δ” indicates the magnitude of the difference in fluorescence between MFC and LFC at the end of the sampling period. (**F**) Quantification of the average difference (δ) in normalized calcium activity between MFC and LFC at the end of sampling across fast (impulsive), typical, and slow (deliberate) strategies. Error bars represent SEM. This difference systematically increases from fast to slow strategies.

To identify the contribution of the different cortical areas, we used a selective optogenetic perturbation approach in behaving head-fixed mice expressing genetically encoded ChR2 in all inhibitory neurons of mouse neocortex (VGAT-cre/ChR2; Fig. 2D, left; see methods)(*29*, *30*). Since the time of the animal’s movement (i.e. taking the decision) was unpredictable in the free-sampling paradigm, it was necessary to use a fixed-time paradigm to confine the optogenetic inhibition to the entire sampling period without overlapping with movement initiation (fig. S6A). We then applied bilateral optogenetic inhibition to distinct cortical regions during a 1-second fixed sampling period (Fig. 2D, left panel; primary visual cortex [V1], n=16 sessions from 7 mice; posterior parietal cortex [PPC], n=12 sessions from 6 mice; retrosplenial cortex [RSP], n=9 sessions from 3 mice; medial frontal cortex [MFC], n=17 sessions from 7 mice; lateral frontal cortex [LFC], n=21 sessions from 7 mice). Optogenetic silencing led to significant drops in performance, primarily in frontal cortical regions, with the largest effects observed in MFC (28.90%±4.5 SEM) and LFC (21.53% ±6.27% SEM) (Fig. 2D, right panel; fig. S6B-F; two-tailed permutation t-test; see methods). These findings are consistent with many previous studies showing that proper recruitment of MFC and LFC over time is necessary for decision-making(*31*, *32*).

To investigate this further, we focused on the flow of calcium activity in the medial-lateral axis of the frontal cortex (MFC→LFC) using the free-sampling paradigm (Fig. 2E). We examined the spatiotemporal dynamics of the frontal calcium activity during the different decision-making strategies, i.e. fast, typical and slow (Fig. 2E, n=6 mice, 18 sessions). Across the spectrum from fast to slow strategies, frontal dynamics showed increasing divergence between medial and lateral regions toward the end of the sampling period (Fig. 2E&F; end). During typical and slow strategies, fluorescence activity in MFC was antipodal (mirror-image) to activity in LFC, despite both contributing equally to performance (Fig. 2D). Notably, LFC was most active just before the decision, when MFC activity reached a minimum. This minimum was most pronounced in slow trials whereas MFC activity at the end of fast trials resembled aborted decisions. Overall, the degree of MFC-LFC antipodal activation at the end of sampling reflected behavioral strategy (Fig. 2F; fast end=0.39 ±0.06, typical end=0.66 ±0.05, slow end=0.73 ±0.04).

### Consistent neural sequence activation during fast and slow behavior

These data imply that activity in MFC at the beginning of the sampling period contributes more to the eventual decision than LFC, for which the crucial activity occurs at the end of the sampling period(*33*). To test this hypothesis, we alternately inhibited both MFC and LFC bilaterally either at the beginning or the end of the sampling period(*34*, *35*) (Fig. 3A). Note that since the timing of inhibition was crucial, it was necessary to use fixed sampling to selectively block either the first (0-0.35s) or last third (0.65-1.0s) of the sampling phase (Fig. 3B). We also repeated the widefield imaging experiments to confirm that the spatiotemporal activity pattern across frontal cortex (MFC→LFC) was consistent under fixed (1s) sampling (n=4 mice, sessions=12; Fig. 3B and fig. S7A&B). In accordance with our hypothesis, optogenetic inhibition in the first third of sampling had a more potent effect when applied to MFC (n=7, sessions=19,16.31% ±2.38% SEM) than LFC (n=10, sessions=19, 8.79% ±2.56% SEM) (Fig. 3C; fig. S7C&D; Early inhibition; see methods). However surprisingly, optogenetic inhibition at the end of sampling impaired performance to a similar degree in both MFC (n=7, sessions=20, 15.18% ±3.20% SEM) and LFC (n=9, sessions=22, 14.37% ±3.37% SEM; Fig. 3C; fig. S7E&F; Late inhibition; see methods), showing that the decay in MFC activity does not result in a decline in its influence on the decision.

**Fig. 3.**
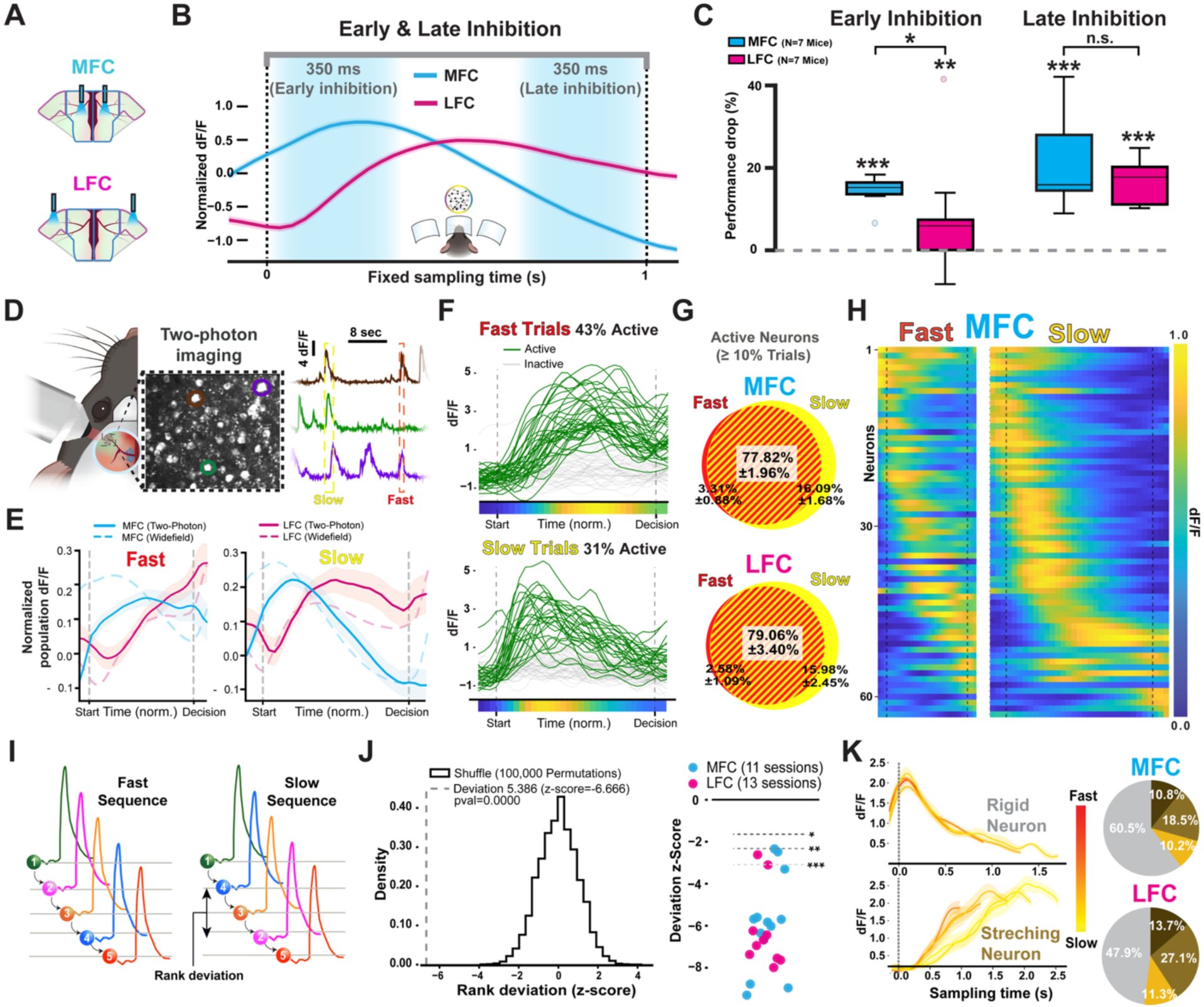
Optogenetic inhibition and two-photon imaging reveal distinct temporal dynamics and neuronal activation sequences in MFC and LFC during fast and slow decisions. (**A**) Schematic depicting bilateral optogenetic inhibition of medial frontal cortex (MFC, blue) and lateral frontal cortex (LFC, magenta). (**B**) Top: Experimental design for targeted optogenetic inhibition in two distinct time windows (early: 0–350 ms; late: 650– 1000 ms) during a fixed 1-s sampling time. Bottom: Average frontal widefield calcium fluorescence during 1-s fixed sampling time (n=4 mice; MFC in blue, LFC in magenta). Shaded intervals highlight early and late inhibition windows. (**C**) Behavioral effect (% performance drop) of early (left) and late (right) optogenetic inhibition of MFC (blue) and LFC (magenta). Significant effects indicated (*p < 0.05; **p < 0.01; ***p < 0.001; n.s., not significant). (**D**) Left: Schematic of two-photon imaging setup targeting calcium activity of layer 2/3 neurons in MFC and LFC. Middle: Example micrograph showing neurons in the imaged cortical area. Right: Example calcium activity traces (dF/F) from three selected neurons (color-coded) highlighting fast (red dotted frame) and slow (yellow dotted frame) trials. (**E**) Mean normalized population calcium activity from layer 2/3 neurons (solid lines) and corresponding widefield signals overlayed (dashed lines) in MFC (blue) and LFC (magenta) during fast (left) and slow (right) behavioral strategies. Shaded areas represent SEM. (**F**) Single-trial neuronal calcium activity traces illustrating active (green) and inactive (gray) trials during fast (top) and slow (bottom) trials. Mini-heatmap reflects neuron’s temporal peak of average activity. (**G**) Venn diagrams showing proportions of active (≥10% active trials) neurons selective for fast (red), slow (yellow), or both behavioral strategies (red/yellow dashed area), in MFC (top) and LFC (bottom). (**H**) Heatmap displaying neuronal population activity during fast (left) and slow (right) strategies from an example MFC session, normalized and ordered by neuronal activation peak during all trials. (**I**) Schematic illustrating the method used to quantify deviations in the neuronal activation sequence (temporal rank) between fast and slow strategies. (**J**) Left: Permuted behavioral strategies rank deviation null distribution (histogram) versus observed rank deviation (vertical dashed line) in an example LFC session. Right: Scatter plot shows rank deviation (z-scored) for neuronal sequences in individual imaging sessions of MFC (blue dots, n = 11 sessions) and LFC (magenta dots, n = 13 sessions). Significant deviations from chance are indicated (*p < 0.05; **p < 0.01; ***p < 0.001). (**K**) Left: Examples of neuron classification based on temporal dynamics. An example of a “rigid” neuron (top) shows no significant correlation (r² < 0.3) of activity peak timing or area under the curve (AUC) with sampling duration. An example of a “stretching” neuron (bottom) with significantly positive correlation (r² > 0.3) between sampling duration and both peak timing and AUC (see methods). Right: Pie charts summarizing the proportion of rigid (grey) vs. stretching neurons in MFC (top) and LFC (bottom). Stretching neurons are defined as those with a positive correlation (r² > 0.3) between sampling duration and either neural activity peak timing (brown), AUC (orange), or both (gold) (see Methods).

The unexpectedly strong influence of MFC at the end of sampling, despite declining widefield activity, raised the possibility that averaging signals across many neurons obscured critical dynamics at the single-cell level. To address this, we used two-photon microscopy to measure the activity of individual neurons directly in MFC and LFC during decision-making (Figure 3D; n=6 mice, sessions=23; 609 active out of 1536 neurons in LFC and 837 active out of 2340 neurons in MFC; see methods). Using this approach, we found that the combined activity of cells recorded using two-photon imaging (Fig. 3E, top) closely reflected the widefield activity (Fig. 3E; fig S8A&B). In particular, the observed decrease in 2-photon activity at the end of sampling correlated with consistently less active neurons in MFC (fig. S8C). This confirmed that the decline seen in the widefield imaging signal was due to fewer active cells which were nevertheless surprisingly critical for decision-making (Fig. 3C, right).Since sampling time differed across trials, we normalized all neuronal activity traces to a common time base separately for fast, typical, and slow strategies, enabling consistent comparison (fig S9A; Fig. 3F).

To characterize neural responses associated with different behavioral strategies, we examined whether fast and slow trials recruit distinct neuronal populations. We found that although the slow strategy engaged a slightly higher proportion of neurons, most neurons were active during both fast and slow trials (Fig. 3G; fig. S9B), allowing for direct comparison of their temporal activation patterns. To do this, we chronologically aligned individual neurons according to their peak activity times to compare activation sequences between fast and slow decision strategies (Fig. 3H; fig S9C). We next tested whether these temporal sequences of neuronal peak activity were statistically consistent across trials, despite substantial differences in sampling duration between fast and slow strategies (Fig. 3H, I; fig. S9C&D)(*36–38*). By comparing to randomly generated permutations, we found similar activation sequences regardless of behavioral strategy (Fig. 3I; fig. S9D). To understand how neurons preserved their sequence rank order, we examined their calcium responses and found that peak activity was stretched or compressed relative to sampling duration(*39–41*) (area under the curve, AUC, and peak time; Fig. 3J bottom; fig. S9E–G; see methods). Neurons with early peak activity (predominantly located in MFC) typically maintained rigid firing patterns (Fig. 3K, top), whereas neurons peaking later often exhibited temporal stretching, a pattern particularly prevalent in LFC (Fig. 3K, bottom; fig. S9G).

These results remained consistent at the population level; averaging calcium activity across all layer 2/3 neurons yielded comparable dynamics at the single cell level to those observed under widefield imaging(*42*) (Fig. S8). Consistently, we also found that the amplitude of population activity at the start of sampling in MFC and not LFC was significantly correlated with sampling duration and easy trial’s performance(*43*) (fig. S10A; fig. S8D). Additionally, MFC neurons showed a tendency to fire sparsely but reliably immediately prior to the decision while LFC exhibited an increasing neuronal engagement with consistent activity levels toward the end of sampling (fig. S10B). In conclusion, optogenetic inhibition and two-photon imaging revealed that the neural mechanisms underlying voluntary decision-making involve both decreasing neural activity in MFC (sparsity) and increasing neuronal engagement in LFC at the time of decision.

### Two modes of operation in MFC versus LFC

To investigate this further, we examined the 2-photon single-neuron responses based on the animal’s choice (Fig. 4A). Interestingly, the pattern of activity during the sampling was very similar for both left and right choices(*37*) (fig S11A). Nevertheless, we identified a subset of neurons as choice-selective, when the peak mean activity during sampling was significantly higher for one choice direction (Fig. 4A, green, “choice-matched”) versus the other (Fig. 4A, brown, “choice-mismatched”). Using this definition, neurons fell into two categories; tuned (either left or right) or non-selective. The peak activity of choice-selective neurons occurred at different times across the whole period of sampling(*44*, *45*) (Fig. 4B; fig S11B). We observed a higher percentage of choice-encoding neurons in LFC versus MFC across all behavioral strategies (Fig, 4C; 33.07 ±3.82%SEM versus 18 ±-3%SEM). However, interestingly, we found that selectivity could vary as a function of behavioral strategy even within a single neuron (Fig. 4D; fig S11C). Next, we asked whether the evolution of choice-selectivity across the sequence of neuronal firing reflected the mode of operation of MFC versus LFC. To quantify this, we binned neurons according to their temporal peak activity and compared matched versus mismatched responses for each bin (Fig. 4E, top panel, slow and bottom panel, fast; MFC 10 session, 182 neurons; LFC 10 sessions, 197 neurons; see methods). For comparison, we also overlaid the average population 2-photon activity from all L2/3 neurons (dotted lines; cf. fig. S8) on the choice-selectivity histograms.

**Fig. 4.**
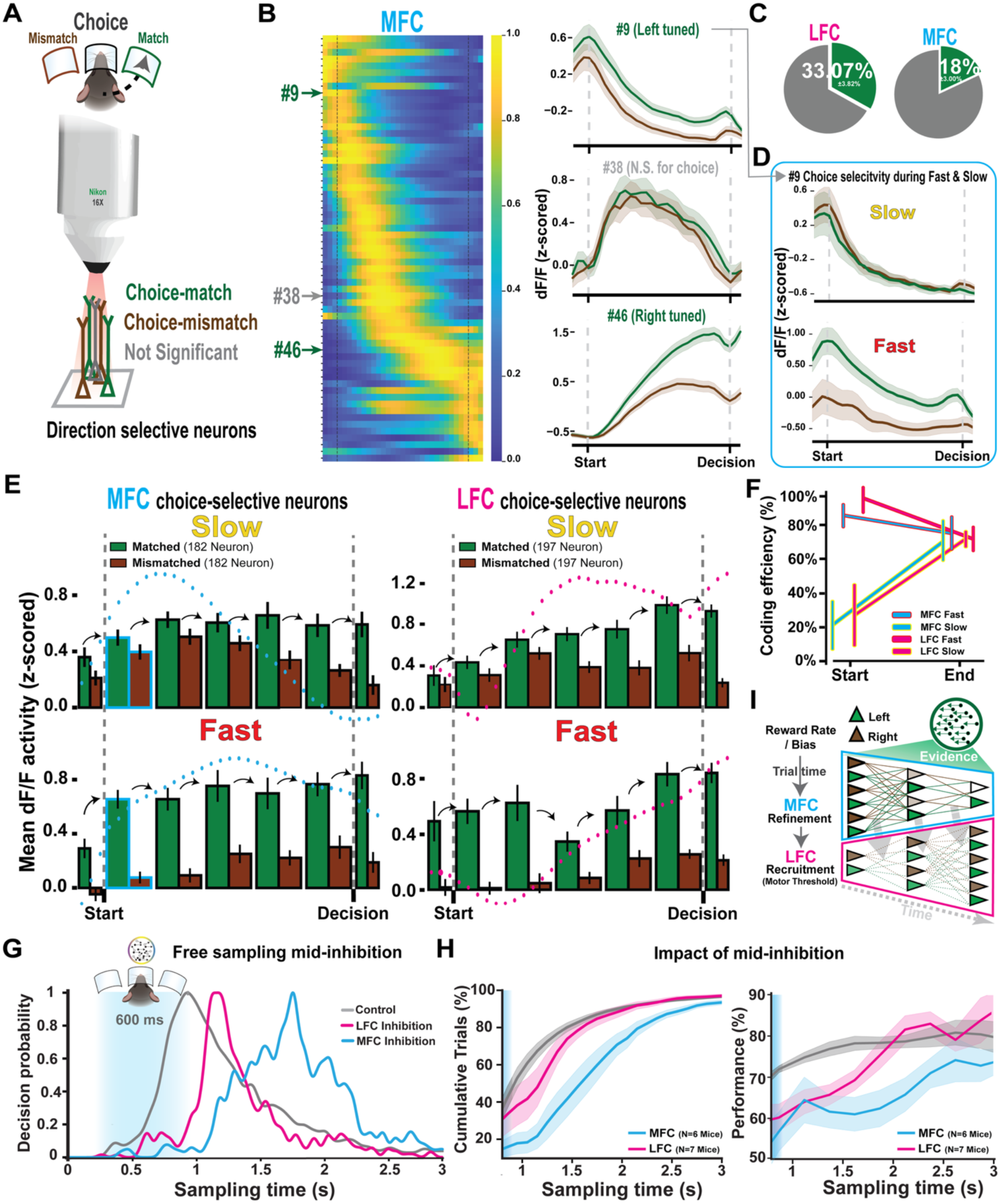
Choice-selective neural sequences in MFC and LFC exhibit distinct modes of operation, with differential contributions to voluntary decision-making. (**A**) Schematic illustrating two-photon calcium imaging of layer 2/3 neurons in MFC and LFC. Neurons are classified as choice-selective based on differential activity patterns during trials in which neuronal preference matches (“choice-match”) or mismatches (“choice-mismatch”) the mouse’s actual choice direction (left or right). (**B**) Left: Heatmap from an example session (MFC) showing the temporal sequence of neuronal activation throughout the sampling period. Right: Three example neurons illustrating activity profiles—selective preference for leftward choice (#9, top), non-selective (#38, middle), and selective preference for rightward choice (#46, bottom). Green/brown traces denote matched/mismatched choices respectively; shading indicates SEM. (**C**) Pie charts quantifying the proportion of choice-selective neurons in MFC (right) and LFC (left). (**D**) Example neuron (#9 from panel B) demonstrating differential responses (choice selectivity) between choice-matched (green) and choice-mismatched (brown) trials during slow (top) and fast (bottom) decision-making strategies. The blue frame indicates the temporal bin of this neuron’s activity in panel E. (**E**) Population responses of sequentially activated choice-selective neurons (ordered by temporal peak, indicated by arrows) for matched (green) and mismatched (brown) trials during correct trials, shown separately for slow (top) and fast (bottom) strategies in MFC (left) and LFC (right). Dotted curves illustrate the corresponding average sum of population activity overlayed. (**F**) Coding efficiency, quantifying the separation between matched and mismatched responses, at sampling start and decision points in MFC (blue) and LFC (magenta) during fast and slow behavioral strategies. (**G**) Top: Experimental design schematic showing timing of bilateral optogenetic inhibition of MFC (blue) or LFC (magenta) during a free-sampling task. Bottom: Example mouse showing probability distributions of sampling durations for control (gray), MFC-inhibited (blue), and LFC-inhibited (magenta) conditions. Shaded region indicates inhibition period (600 ms). (**H**) Left: Cumulative distribution of sampling durations across all mice for control (gray), MFC inhibition (blue), and LFC inhibition (magenta). Right: Behavioral performance (percentage correct) following the offset of inhibition across the sampling time, comparing control, MFC-inhibited, and LFC-inhibited trials. (**I**) Summary schematic illustrating how different parameters (e.g., reward rate, sensory evidence, previous choice) shape decision-making via distinct modes of frontal cortex operation. MFC primarily refines decisions by integrating these parameters gradually, while LFC facilitates commitment to action once evidence crosses a motor threshold, particularly differentiating fast from slow strategies.

We conclude that slow behavior commenced from a more balanced state of choice representation (matched vs mismatched) at the start of sampling, a trend consistent in both MFC and LFC. In contrast, fast behavior was characterized by an initial state of imbalance in matched versus mismatched activity. Additionally, a consistent observation across all conditions was the emergence of an imbalance in choice representation by the end of the sampling period (fig S11D&E).

In general, these results can also be interpreted using a metric we term “coding efficiency,” defined as (matched – mismatched) / matched, reflecting the propensity of neural activity to favor the animal’s chosen direction at any stage in the sequence (Fig. 4F; fig. S11F). This metric allows for quantification of how neuronal activity evolves to influence choice selection. We observed that coding efficiency substantially increased during the sampling period for slow behavioral trials but decreased for fast trials. To further elucidate the neural mechanisms underlying these differential strategies within the DDM+Q+R computational framework (Fig. 1I-K), we found that a substantial fraction of neurons multiplexed signals encoding previous trial outcomes together with representations of previous or current choices (fig. S12A&B). Notably, reward history significantly sharpened neuronal selectivity for both previous and current choice signals during the early phase of sampling (fig. S12C). Given the limited availability of sensory evidence at the early stage of sampling, we interpret this early imbalance representation as a deviation in the starting offset or noise signal during evidence accumulation. This reward-history-driven bias provides a mechanistic link to the fast trial strategy, marked by strong priors and elevated coding efficiency at the start of sampling (Fig. 1I; Fig. 4F; fig S13).

Our data so far suggest that MFC and LFC operate in distinct but complementary modes during decision-making. Both regions exhibit improved choice coding efficiency during slow trials and reduced efficiency during fast trials (Fig. 4F). Yet they appear to achieve this via different mechanisms. In MFC, this improvement is driven by a refinement process in which the number of neurons representing choice decreases more steeply for mismatched activity than for choice-aligned signals (Fig. 4E, left, fig S14). In contrast, the number of active LFC neurons appears to gradually increase over sampling to a consistent threshold value for triggering action (Fig. 4E, right, fig S14). The distinct temporal trajectories in MFC and LFC, despite converging coding efficiency, suggest they may contribute differently to how information is sampled and integrated over time(*5*).

To test this idea, we asked whether interfering with the sequence of firing during the sampling period would differentially affect behavior. Specifically, we inhibited MFC or LFC midway through free sampling (0.3–0.9s), interrupting the sequence and decision-making process (Fig. 4G, blue shaded area). Inhibiting LFC briefly disrupted the decision-making process, reflected in sampling duration and performance, which quickly recovered after the end of inhibition (Fig. 4G-H, magenta; fig. S15). In contrast, inhibiting MFC caused much longer delays in sampling and a sustained drop in performance which did not recover even with long sampling times (Fig. 4G–H, blue; fig. S15). This indicates that decision-making is more sensitive to the sequence of activation of neurons in MFC than LFC.

These findings lead us to revisit the conceptual framework of serial sampling, highlighting the distinct contributions of MFC and LFC to decision-making (Fig. 4I). In this framework, MFC generates a sequential refinement of choice representations, culminating in a small population of neurons encoding the eventual choice (Fig. 3C & Fig. 4G&H). Disrupting this sequence at any point impairs performance in a sustained manner, suggesting that MFC actively tracks the evolving state of evidence. In contrast, LFC shows a rapid rebound following transient inhibition, implying that it does not accumulate information locally but rather monitors upstream signals (possibly including MFC) to determine when to act. This interpretation aligns with recent findings that LFC activity reflects the decision state but may not integrate evidence directly(*46*, *47*).

Taken together, these results suggest that MFC plays a central role in computing what decision to make, while LFC determines when to act. Crucially, our data also imply that the MFC may exert an inhibitory influence over downstream motor initiation, releasing this brake only once its internal refinement process converges. In this framework, disrupting MFC not only degrades choice quality but also delays commitment, whereas disrupting LFC mainly affects the timing of action execution. This division of labor between refinement and commitment offers a mechanistic account of how serial sampling is coordinated across frontal regions and may generalize to other forms of voluntary decision-making.

## Acknowledgement

We thank Moritz Drüke and Arina Lobova from the Larkum Lab for insightful discussions. We are also grateful to all members of the Larkum Lab for their valuable input and to the supporting staff for assistance with organizational tasks. Special thanks to Alexander Schill and Jan Ode from the Research Workshop at Charité – Universitätsmedizin Berlin for developing and manufacturing the experimental devices. We deeply appreciate the support of Dr. Claudia Abramuik and the Charité Mitte animal facility for providing access to facility space and equipment, as well as for their assistance in breeding various mouse lines. Special thanks to Joao Couto for advice on using the widefield preprocessing library.

## Data and materials availability

All data required to evaluate the conclusions of this study are included in the main text and/or the Supplementary Materials. Source data and analysis code will be deposited in a public repository and made available prior to publication.

## Supplementary Materials for

## Materials and Methods

### Human psychophysical Experiment

The study was conducted at Humboldt university and approved by the local ethics committee of the Humboldt University Berlin. 56 students constituted the pool of participating subjects. As students participated on a voluntary basis but varying motivation, only the data of the top 30% best performing subjects of each group cohort were used. Each subject final score was based on their performance in both accuracy and maximum correct outcome experiments (n=18 Subjects). The experiment was conducted on 6 group cohorts, each involving 9-13 subjects. Subjects were assigned a computer rig (Dell Optiplex 7400 AIO Series), were made aware of the scoring criteria and were encouraged to compete for the highest score. Most users ran both accuracy and maximum correct outcome experiments, with each experiment duration set to 30 minutes including a 5-minute break at the 15-minute mark, as well as a mandatory 20-second break every 100 trials. Subjects performed an initial 10 test trials. For the accuracy competition, a subject’s score is calculated as the percentage of correct trials over the total trials, with a minimum requirement of 50 trials. For the maximum correct outcome competition, the subject’s score is the total number of correct trials. A central server controlled by the experimenter activated a countdown timer to synchronize the start of the session on all subjects’ computers, thus ensuring all subjects are participating concurrently. Subjects received correct/incorrect feedback after each trial and could track their ranking among other players, viewing the leaderboard during individual 20-second breaks and the 15-minute group break. The experiments were implemented using jsPsych: and utilizing the jspsych-rdk plugin(*48*, *49*). The stimulus comprised of 1,000 white dots on a black background, with a dotliftime of 1 second and a dot size of 0.5 pixels viewed by subjects at≈50cm distance resulting in a dot size≈0.02°.

### Mice Experiments

All experiments were conducted in accordance with the guideline given by Landesamt für Gesundheit und Soziales Berlin and were approved by this authority.

### Surgical Preparation

We used adult (>P40) C57BL/6 wild-type mice, GP4.3 thy1, (#:024275, The Jackson laboratory), and VGAT-CHR2-EYE transgenic mice (#:014548, The Jackson laboratory). Mice were anaesthetized with mit Ketamin/Xylazin (Ketamin 12,5 mg/ml, Xylazin 1 mg/ml (10 µl/g) intraperitoneal) and kept on a thermal blanket. After surgery,mice received Caprofen (5 mg/kg) und Buprenorphin (0,05-0,1 mg/kg) for 2 days. During head-fixation skin and periosteum were removed, a thin layer of light-curing adhesives was applied to the skull. A light-weight head-post was fixed on the skull with a dental cement. For widefield imaging, intact skull cleared using a layer of cyanoacrylate (Zap-A-Gap CA+, Pacer technology) to clear the bone(*50*). Green fluorescent was check under widefield microscope after surgery and double checked later with two-photon imaging calcium imaging. For optogenetics, over intact skull stereotaxic marks were identified using permanent marker, before skull clearing, to label different targets of optogenetic inhibtion: primary visual cortex, V1 (1.0 mm anterior to lambda and 3 mm lateral from the midline), retrosplenial cortex, RSC (1.0 mm anterior to lambda and 0.7 mm lateral from the midline), posterior parietal cortex, PPC (2.0 mm anterior to lambda and 2 mm lateral from the midline), Medial Frontal Cortex, MFC(0.8 mm anterior to bregma and 0.7 mm lateral from the midline), LFC, LFC (2 mm anterior to lambda and 1.8 mm lateral from the midline). For two photon imaging, a 3-mm craniotomy was made 0.3mm radially anterior from bregma and sealed with a 3-mm glass coverslip with cyanoacrylate glue and dental cement.

### Behavior Behavioral Setup

Our behavioural studies employed a modification for the setup developed by Nashaat et al. 2016, that allows integrating natural behaviors with advanced neuroimaging in a head-fixed context. The setup is composed of a Plexiglas air-table, perforated with evenly spaced holes, rerdirecting compressed air to generate an upward current flow, levviating a PLA platform (15cm diameter) on top. The platform features 3 nose-pokes allowing the mice to navigate and seek rewards. A head-fixation mechanism ensures the mouse’s head remains centered. The nose pokes, equipped with LEDs, infrared sensors, and a reward delivery tube, controlled by a solenoid valve, are extension of the Bpod (Sanworks) behavioral control system. To display RDK stimuli, a Retina LCD screen was positioned in front of the platform, approximately 10-15 cm away. RDK stimulus was generated with custom script utilizing either Psychtoolbox or Psychopy(*51*, *52*). White dots were rendered against a black background. A single dot is set to 2.5° size with a lifetime of 0.5s and travelling speed of 20°/s-40°/s, with wide-field sessions and 2-Photons sessions always using 20°/s. Dots were rendered to fill the whole screen with dots filling 20% of viewing apreature. Dots traveled in 8 directions (cardinal- and inter-cardinal directions) with a ratio of dots, determined by the trial’s coherence level, formed a majority moving either left or right, toward the direction of the reward port.

### Behavioral Training

The initial phase introduced freely moving mice to the behavioral system using a simple light-chasing task conducted over three sessions, each approximately 30 minutes. This phase allowed the mice to acclimate to the environment and task structure. In this task, trials began with the mice fixing their nose inside the central port, illuminated first, followed by a light cue on either the left or right port signaling the correct choice. A successful selection resulted in a water-based reward, sweetened with 0.05% saccharine and sucralose, while incorrect choices triggered a high-pitched beep (4000-5000 Hz) as negative feedback. Subsequently, mice were head-fixed to adapt to moving the platform themselves, guided by experimenter-assisted platform movements. As mice perform above 75% accuracy in light-chasing task, a 4-seconds timeout penalty for incorrect choices is introduced. Trials where mice exited the central port before minimum sampling resulted in aborted trial with 1-2 second timeout. Expert-level was marked by sustained >75% accuracy across sessions, after which RDK stimuli were gradually integrated alongside light cues. Initially paired, the reliance on light cues was reduced until the task relied solely on RDK stimuli, with varying coherence levels (100%, 50%, 40%, 10%, 0%) introduced as the training advanced. Motion direction and coherence varied pseudo-randomly to prevent biases.

### Fixed sampling Paradigm

For the fixed time (FT) Paradigm, the approach mirrored tasks previously conducted(*10*, *11*, *53*). In FT trials, Mice must remain poked-in for the entire 1s stimulus duration. Early-Withdrawals counted as aborted trials. For the chronometry experiment (Extended Fig. 1c), trials between 0.3s to 1.2s, in increments of 0.3s, were presented in pseudo-random order with 20% probability in-between 1s trials.

### Free sampling Paradigm

For free sampling paradigm, the approach mirrored tasks previously conducted with humans and monkeys. In this paradigm, the stimulus is continously presented and only terminated on exiting the center port, with a maximum of 5s cap. The mice had the autonomy to end the sampling period anytime between a minimum of 0.3s and a maximum of 5s, facilitating a balance between impulsivity control and decision-making flexibility. Trials with less than 0.3s sampling duration are counted as aborted trials.

### Optogenetics

The targeting coordinates for optogenetic inactivation were determined using the Allen Brain Map API for Mouse Brain data(*29*, *35*). 473-nm laser (OEM Laser Systems) was administered through bifurcated fiber bundle (Thorlabs BFYL2LF01), each emitting light covering approximately 250 μm in diameter at a 40 Hz with an 80% duty cycle and an intensity of 6 mW/mm² to ensure that photoinhibition was confined to the targeted cortical areas(*29*). Additionally, blue masking light was positioned above the viewing screen. Training data indicated that mice exposed only to the masking light maintained an average performance exceeding 75%. In each experimental session, mice experienced perturbation in one brain area per session, with optogenetic trials pseudorandom introduced among control trials. To sustain high performance levels (70% or higher) and avoid demotivation, the probability of light illumination was set at 15-20%. During optogenetic inactivtion under fixed-time paradigm, inactivation duration was either throughout the entire delivery period of sampling (1s, Full), during early sampling (0-0.35s, Early), or the late sampling (0.65-1s, Late) phases of stimulus presentation (Figure 2)(*34*).

### Widefield imaging

We used an adaptation of widefield imaging setup that utilizes two light source for hemodynamic responses correction. We used the following variation: for the inverted tandem lens macroscope, we utilized a top lense with focal length, 105 mm (DC-Nikkor, Nikon) and a bottom one with focal length 85 mm (85M-S, Rokinon)(*28*, *50*).

### Two-Photon Calcium Imaging

Imaging utilized a resonant-scanning two-photon microscope (ThorLabs B-scope) at a 5 kHz scanning rate. GCaMP6s excitation occurred at 940 nm via a Ti: Deepsee laser (90-110 mw power). Data collection was through a 16x water immersion objective (Nikon, 0.8 NA), capturing 512 x 512 pixel full-frame images of layer II/III neurons within a 1 mm² area, at 150-300 µm depths. Selection criteria for imaging fields included cell density and minimal vascular interference. Imaging operated at 30 Hz frame rate, employing ThorImage software.

### Data Analysis

Data analysis was performed using Python utilizing Numpy, Scipy, Pandas Scikit-Learn, Matplotlib, Seaborn and statsmodels libraries.

### Human and Mice Behavioral Analysis

Each session trials were grouped by difficulty. Within each difficulty, the data were grouped into 3 thirds (1^st^ Quantile: Fast, 2^nd^ Quantile: Typical, 3^rd^ Quantile: Slow). In mice, early-withdrawal trials, i.e. trials where mouse spent <0.3s sampling (88 trials), and trials where mice stayed in the center-poke after the end of the maximum 5 seconds stimulus presentation were excluded (131 trials). Additionally, mice with few trials (<300 trials) or that followed a non-typical strategy, dissimilar to Fig. S2 E, where correct trials are slower than incorrect trials, were excluded (21,628 out of 90,973 Trials were excluded). For human subjects, stimulus was set to be indefinitely presented. Here, trials with more than 5 seconds were excluded (0.2% of maximum correct outcome competition, 7.84% for accuracy competition).

In analyses where reaction-time Z-score normalization is applied (Fig 1D, G-H, Sup. Fig. 3. A-B,E-G) each mouse’s data is pooled across all its sessions. If only a subset of data is selected, e.g. only correct trials in Fig 1G, then Z-score normalization is first applied beforehand. For human data, for analyses that characterize each experiment type (Fig. 1F-H. Sup. Fig. S2 G-H,J-K), data was normalized only for the given experiment type. For analyses characterizing change in behavioral signature across experiment types (Fig. 1C, Sup. Fig. S2 B-C) data was normalized across both experiment types.

Psychometric curves (Fig. 1 E-F, J Right, Sup. Fig. 1 B Middle, Sup. Fig. 2F,I, Sup. Fig. 4 A, Sup. Fig. 6 B-F, Sup. Fig. 7. C-F, Sup. Fig. 15 E-F) were fit using the Psychofit library (https://github.com/cortex-lab/psychofit) using error function configuration. Single-sided psychometric functions, characterizing performance and perception threshold, were fit using a single-lapse-rate configuration, while two sided psychometric functions, additionally characterizing choice bias, were fit using two-lapse-rate configuration. As the region of greatest rate of change within the psychometrics differ between humans and mice subjects, human data were binned in [0, 1, 5, 10, 20, 50, 100] bin intervals. Mice data were binned in power of two increments, yet when excluding the empty bins the remaining yield [0, 16, 32, 64, 100] bin intervals. For Fig 1I, Sup. Fig. 2L and Sup. Fig 3I, a paired t-test was performed across each subject.

For Sup. Fig. 2B-C, mice were limited to a maximum sampling time of 3 seconds to match the maximum stimulus time used in human experiments. To compare the 3 groups, and after testing each group for normality (Shapiro-Wilk test), we deployed one-way ANOVA followed by Tukey’s test with Bonferroni correction for post-hoc pairwise comparisons.

In Sup. Fig. 3F, groups were compared using the Kruskal-Wallis test, with post-hoc comparisons performed using Dunn’s test with Holm–Bonferroni correction. In Sup. Fig. 3G, trials were pooled acris all mice and were classified as Win (previous trial correct) or Lose (previous trial incorrect). Each classification was split into bins of 800 trials each, where the stay/switch update value within each bin was calculated relative to the system-generated baseline stay/switch distribution within the given bin. A Gaussian kernel was applied to smooth changes between neighboring bins. Two mice were excluded due to abnormal low z-score distribution (mean percent of trials < −1 z-score=3.28%, excluded mice=18.62% & 13.26%). The stay/switch update per sub-panel in Sup. Fig. 3H was calculated similarly to a single bin calculation in Sup. Fig. 3G but without Gaussian kernel smoothing.

Average Reward-Rate figures (Fig 1H, Fig. S3 B-D) are calculated per subject then subjects’ mean of means is calculated. A subject average reward rate (including Fig 1K, Fig. S4 Reward Rate column) is calculated as the subject’s sessions’ mean of means sampling for each reward values over the last 5 trials. A session’s average reward rate entry with 5 trials or less is excluded. As Avg. reward rate value=0, i.e. no reward over the last 5 trials, is the least occurring event with few data points, it is excluded from the plots.

To quantify the contribution of individual predictors to sampling time (Fig. S3 J), we used ordinary least squares (OLS) regression from the statsmodels library, with the model log-likelihood serving as the measure of fit. For each animal, we log-transformed the sampling times and fit both a null model (intercept-only) and a full model including all predictors. The difference in log-likelihood between the full and null models was used to define the total log-likelihood improvement — a measure of the overall explanatory power of the predictors. To assess the unique contribution of each predictor, we fit reduced models omitting one predictor at a time and computed the percentage loss in log-likelihood improvement relative to the full model. This procedure was repeated for each subject, and group-level statistics were computed as the mean and standard error of the mean (SEM) across animals. (Mention here the parameters used, and they were used to avoid redundancy information between variables)

### Model

To model the effects of incorporating Q-learning and R-learning reinforcement learning influence on the Drift Diffusion Model (DDM) (Fig. 1J-L, Fig S4), we fitted each mouse’s behavioral data against four models:

### DDM

We used the standard DDM as the baseline comparison model: Let offset starting point:

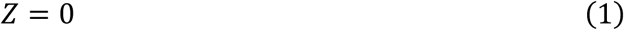

Let the initial evidence state be:

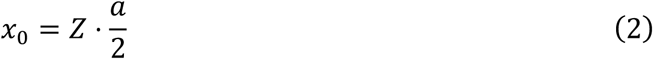

The decision variable evolves according to:

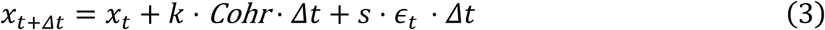

A decision is made when *x_k_* reaches either bound (± a/2). The time to decision is:

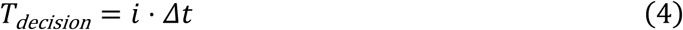

Trial Total decision time is calculated as follows:

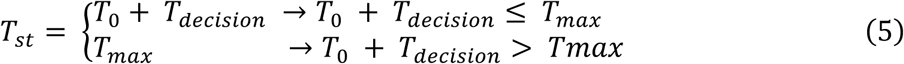

Where *a* is the distance between decision bounds at ± a/2, (*a* > 0), *Z* is normalized starting point (*Z* ∈ [−1,1]). *x_t_* is accumulated evidence at time step *t*. *Δt* is a time step increment (Δ*t* > 0). ***Cohr*** is the dots coherence and direction on the trial (*Cohr* ∈ [−1,1]). ∈_t_ is random standard normal noise at time step *t* (∈_t_ ∼ *N*(0,1)). *T***_decision_** is time until a boundary is crossed. *T_0_* is number of time steps until boundary crossing. *T*_0_ is non-decision time, e.g., motor and sensory latency. *T****_max_*** is maximum simulation time per trial. *T***_st_** is trial total time including non-decision time.

Model free parameters are: *k*: drift scaling factor (*k* > 0), *s*: noise scaling factor (*s* > 0).

### Q-learning + DDM

To capture action preferences, we modeled a simple Q-learning process starting from a center nose-poke state *S*_0_, with left and right nose-pokes as terminal states each associated with an action value. We allowed an additional parameter to capture motor bias:

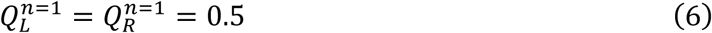

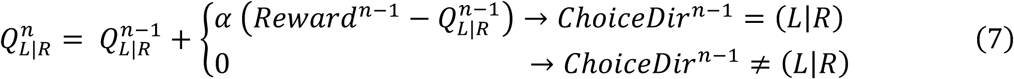

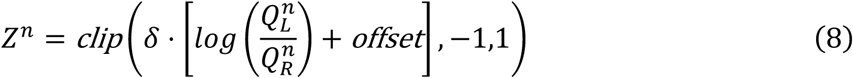

Where *n* is the trial number within a session. ***Reward*** ^*n*^ flags whether a reward is received on trial *n* (*Reward* ^n^ ∈ {0,1}). ***ChoiceDir*** ^*n*^ is choice direction on trial *n* (*ChoiceDir* ^n^ ∈ {0,1, *nil*}). *Q*^*n*^_L_, *Q*^*n*^_R_ are action values for left/right at trial *n* (*Q*^n^_L|R_ ∈ [0, 1]).

Model free parameters are: ***offset***: motor bias term (*offset* ∈ [−1,1]), α: Q-learning update rate (α ∈ [0,1]), δ: starting offset scaling factor (δ ∈ [0,1]).

### R-Learning + DDM

To model adaptive strategy changes in response to reward rate, we applied R-learning as modulating of noise gain rather than a change in decision bound. This choice was guided by the latter findings of the constant motor threshold value of LFC.

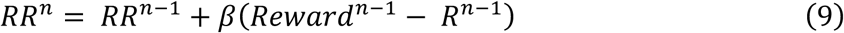

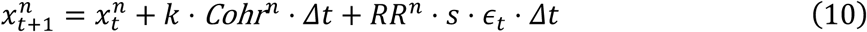

Where *RR*^*n*^: reward rate at trial *n* (*RR*^n^ ∈ [0,1]).

Model free parameters: β: R-learning update rate (β ∈ [0,1]).

### Q-Learning + R-Learning + DDM

We combined both Q- and R-learning models to evaluate their joint influence on behavioral strategies. In all models, to reduce interaction effects when fitting both the decision boundary and noise parameters simultaneously, we fixed the decision boundary to *a* = 2, setting bounds at (−1, 1). Using differential evolution optimization, models’ loss functions were minimized using Chi-Square fitting method to lessen the difference between the subjects’ sampling times and those generated by the model(*54*). Loss was first computed separately for correct and incorrect choices, and the resulting loss values were then summed. For each choice condition, the Chi-square statistic was calculated using the following quantile intervals from the subject’s data: [0, 0.1, 0.3, 0.5, 0.7, 0.9, 1]. The integration step was set to Δ*t* = 0.005s, with a maximum allowable sampling time of *T*_max_ = 3*s*. Trials that did not reach a decision threshold within this time were categorized as no-choice and no-reward trials.

The coefficient of determination R² (Fig 1L) is computed for both the 2-sided psychometric, thus capturing performance and choice bias, and the average reward rate. For psychometric R², the difference in each difficulty-direction bin between slow and fast psychometric fits of the subject’s real data is compared against the corresponding data in the model. For reward-rate R², sessions’ average reward-rate for each reward-rate value is compared in real-data vs model data. A session’s reward-rate value entry with less than 5 trials is dropped.

### Optogenetics Data Analysis

Optogenetics data was collected for one brain-region at a time and its control trials span between the first and last brain-region optogenetics trials. To quantify the optogenetics performance effect, we computed the drop in performance across each session as follows:

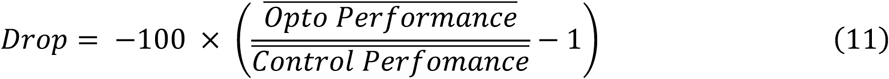

We adapted the approach in to test whether optogenetics inhibition effect is significant (Fig. 2D)(*29*, *30*), data were pooled across mice, and for each of 10,000 iterations, subjects, sessions, and trials were resampled with replacement in that sequence. For each iteration, change in performance was computed. The p-value was determined by calculating the proportion of iterations where the performance changed sign, e.g. drop in performance changed to performance improvement, was detected. As a two-tailed test was used, the significance threshold was set at 0.025 and adjusted for multiple comparisons using the Holm-Bonferroni correction.

To compare whether the performance effect of optogenetics inhibition significantly differs between two brain regions (Fig. 3C), we first perform Shapiro-Wilk test for normality on both groups. If data is not normally distributed for any of the groups (Fig 3C left), Wilcoxon–Mann–Whitney two-sided test is used, when data is normal for both groups (Fig 3C right), two-sided t-test is used.

To regress the relation between sampling time and performance under control and optogenetics inhibition conditions (fig. S15C), we performed linear regression on the z-score normalization on the sampling time of each animal control session. The optogenetics trials were z-score transformed using the corresponding control data distribution. To acquire mean performance, trials were grouped in bins of 50 trials. A linear fit for each of the control and optogenetics sessions as well as the overall data were plotted using a fixed common origin value at −1.5 z-score and 50% performance.

### Wide-Field Imaging Data Analysis

We adapted a variation of the analysis in (https://github.com/jcouto/wfield) with the following changes: 1) using 800 singular value decomposition (SVD) components instead of default value (200 components) to reduce dimensionality-reduction loss. 2) Following image registration step, pixels outside the brain were set to black to minimize their assigned SVD weights. 3) we used separate left and right hemispheres for a better fit that accounts for unequal sizes due to zooming artifacts caused by small angle tilt in the acquisition camera. To characterize the wide-field activity of a typical trial (Fig. 2B, left), we constructed a reference set by pooling median-time trials across sessions (fig. S5B). For each session, we identified trials falling between the 0.45 and 0.55 quantiles of the sampling duration distribution (fig. S5B, vertical colored bars). We then computed the global median of these session-level medians (fig. S5B, red horizontal line). The reference set included all trials whose sampling durations were within ±0.1 seconds of this global median (i.e., vertical colored bars falling between the horizontal dotted lines).

### Two-Photon Data Analysis Preprocessing

Raw two-photon imaging data were preprocessed using Suite2p for image registration and neurons extraction followed by manual inspection(*55*). To identify active neurons and neurons active trials, we adapted a variation of the steps used in previous 2 photon imaging studies(*44*). We extracted raw traces 0.1 seconds before start of sampling until 0.1 seconds after end of sampling. For each trace, a z-score normalization is temporarily applied across a concatenation of all the sampling trials. Each trial is re-extracted, and the standard deviation is computed after applying a 1-frame gaussian smoothing. A neuron is marked to have an active trial if the standard deviation of the trace within the trial exceeds a threshold value of 3 x 5^th^ percentile of all the trials standard deviation. Neurons with less than 5 percent active trials are discarded. 837 out of 2,340 neurons in MFC (36.44% ±6.9%SD per session), and 609 out of 1,0536 neurons were marked as active in LFC (38.88% ±7.51%SD per session). Neurons ΔF/F_0_ was calculated using the whole session trace after applying a gaussian smoothing with a standard deviation=1. F_0_ is computed by computing the mode value of all the frames baseline. A frame’s baseline was computed as the maximum of the minimum values in the surrounding 60 seconds’ window.

### Heatmaps

Fast and Slow heatmaps (Fig 3H) were sorted in reference to all trials’ heatmap or in reference to all Slow trials (Fig S11 B). The activity is normalized across the minimum and maximum values found in both heatmaps.

### Tuning selectivity

To test neuronal selectivity for specific criteria, e.g left/right choices (Fig. 4C) and previous left/right choices (Fig. S12 A, we extracted the maximum trial activity for each condition and compared them using a two-sided Wilcoxon–Mann–Whitney test.

To quantify the percentage of neurons that are modulated by previous trial reward outcome (Fig. S12 C Pie-Charts), we computed a modulation index for each neuron, defined as the difference in activity between matched and mismatched stimulus conditions, separately for trials following a correct and incorrect outcome. We then calculated the change in this modulation index based on the previous trial’s outcome. To assess statistical significance, we performed a permutation test of 1,000 shuffles in which trial labels for previous outcome were shuffled, generating a null distribution of selectivity differences.

To determine the percentage of neurons sensitive to reward history (fig. S12B), we binned each neuron’s trials based on reward history, capping the history at a maximum of three consecutive correct or incorrect outcomes. If the neuron was determined to be signficantly selectivy tuned to a criteria (e.g. previous-choice left/right), the analysis is then restricted to trials matching neuron’s selectivity (e.g preferred direction), otherwise all the trials are used. We computed the mean activity for each bin and ordered bins in descending sequence of incorrect choices followed by ascending correct choices (aligned with the reward history timeline in fig. S12C, top). Neurons whose activity increased monotonically across this sequence were classified as reward-history sensitive. These results were compared to a null distribution generated by 1,000 shuffles of reward-history labels across trials.

### Sequence characterization

To determine each neuron’s rank within a sequence, we used the median peak firing position across its active trials within time normalized trials.

To assess the consistency of sequential neuronal firing across trials (Fig. 3I–J), we compared the rank deviations between fast and slow trials. For each session, a neuron’s rank was defined as its median peak firing position in fast and slow trials separately. Tied ranks were assigned the same value, with subsequent indices skipped (e.g., 1, 2, 2, 2, 5). The rank deviation between fast and slow trials was calculated as:

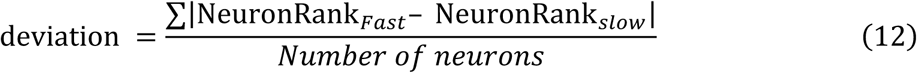

We then performed a permutation test with 100,000 iterations, shuffling the ranks to generate a null distribution. The original deviation’s p-value was projected onto the resulting z-score distribution. A variation of this analysis (fig. S9D–E) was also conducted, comparing fast and slow trial ranks to those from typical trials (defined as the middle third quantile). Permutation tests were run by reshuffling the typical trial ranks. This approach allowed us to assess whether the typical sequence of activation resembled those observed in both fast and slow trials, using an unbiased, independently derived reference set for evaluating rank consistency.

To investigate how single neurons modulate their responses with respect to sampling duration (Fig. 3K, fig. S9F–H), we analyzed the correlation between each neuron’s firing position and area under the curve (AUC) of calcium activity as a function of increasing sampling time. AUC was computed from calcium traces while accounting for two key caveats. First, strong motion artifacts in the two-photon imaging field, particularly along the Z-axis, can induce sharp frame-to-frame deflections in the calcium trace that are inconsistent with expected GCaMP6s dynamics. While such artifacts are typically mitigated in trial-averaged data, single-trial traces remain more sensitive. To reduce these effects, we applied Gaussian smoothing with a standard deviation of 2. The resulting traces (Fig. 3K left; fig. S9F,G left panels) were visually inspected to ensure removal of high-frequency artifacts. Second, the slow decay kinetics of GCaMP6s fluorescence can cause residual signal from prior events to appear in subsequent time points. To ensure AUC reflects ongoing or sustained activity, we restricted calculations to positive or sustained deflections. A trace may contain zero or more such events. For each, AUC was defined as the sum of trace values above 60% of the peak amplitude within the event window (i.e., ≥ 0.6 × [peak – value at deflection onset]). Using neurons’ active trials (see “2-Photon Preprocessing” for the definition), we performed linear regression of both AUC and peak firing position against sampling time, and computed Pearson correlation coefficients (fig. S9F,G middle and right panels). To avoid skewing results, trials with sampling durations ≥ 3 standard deviations from the mean (Z-score) were excluded. Neurons with an absolute Pearson correlation ≥ 0.3 were considered non-rigid (Fig. 3K right; fig. S9H).

To further assess the significance of these correlations, we plotted histograms of observed values against distributions obtained via 1,000 reshuffled controls, in which sampling times were randomly permuted within each neuron (fig. S9H). Notably, neurons with early peak firing positions can exhibit gain modulation (i.e., changes in AUC), but their firing time remains fixed (fig. S9H top). In contrast, neurons peaking later in the sampling epoch can exhibit both gain and timing modulation (fig. S9H middle and bottom).

In Fig.4E and fig. S11C, each pair of bins contain the set of significantly direction-selective neurons, pooled across all brain-region sessions, that their peak firing position IQR fall within the bin-pair boundary. Thus, each pair of green and brown bins contain the same set of neurons under different trial conditions: when the trial matches the neuron selectivity (green) and when trial mismatches it (brown). The bin height and error bar reflect the average and SEM of the neurons mean z-score activity under the bin condition. We split time-normalized sampling duration into 7 bins pairs: boundaries pair before and after sampling start/end and 5 equal-time pairs during sampling. We excluded neurons that their median peak firing position falls within the first and last bins pairs, to avoid including predominantly motor tuned neurons.

To further confirm the observed effects, a logistic-regression choice classifier (Fig S11D) was run on significant choice neurons. After applying z-score normalization to each significant neuron traces during the sampling epoch, the features input to the classifier is each significant neuron maximum firing at each trial at either the start of sampling (0.1 before and 0.2 after start of sampling) or end of sampling (0.2 before and 0.1 after end of sampling). The classifier input labels are the animal direction choice. For each test combination, a 1,000 of random training and test splitting is performed, with 30% of the data used for testing on each run.

We defined Coding-efficiency (Fig. 4F), a measure of the mean separation between match and mismatched selectivity, as follows:

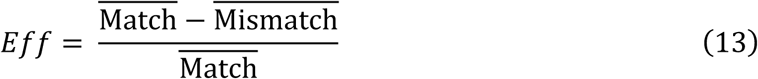

Coding-efficiency error bars are computed using the propagation of uncertainty approximation method:

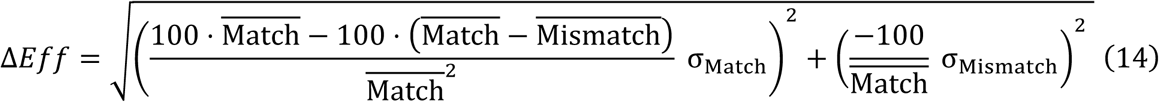

To describe a two-photon population activity, either the mean activity of all neurons (Fig. 3E solid lines, Fig. 4E background dotted lines, Fig. S8), or the mean activity of only significant neurons (Fig S10 A left, B left, Fig S11 C bottom), the sum of z-scored activity is computed across each time point.

To investigate whether early neural activity within a brain region influences fast versus slow behavioral strategies (fig. S10A, middle), we quantified the percentage of active neurons during the early sampling window (from 0.1 seconds before to 0.3 seconds after sampling start), relative to the total number of traces in the session (see “2-Photon Preprocessing” for the definition of active trials). The 0.3-second cutoff corresponds to the minimum enforced sampling duration in the behavioral task. Trials were pooled across animals and binned into 0.25-second intervals based on their sampling durations. A linear regression and Pearson’s correlation coefficient were then computed to quantify the relationship between early activity and sampling duration.

To examine how the density, or sparseness, of population activity within a brain region evolves with sampling duration (fig. S10B, right), we repeated the early activity analysis described above (fig. S10A, middle), but applied it to the final phase of the sampling period. Specifically, we analyzed the window spanning the last 30% of the sampling duration up to 0.1 seconds after the end of sampling (i.e., movement start).

To investigate whether early neural activity within a brain region influences performance (fig. S10A, right), we repeated the early activity analysis described above (fig. S10A, middle) with the following modifications: only easy trials were included to control for task difficulty, and trials were binned by early activity levels, ranging from 0% to 30% in 3% intervals. For each bin within a session, performance was calculated as the difference between the mean performance of trials in the bin and the mean performance of all easy trials. A linear regression and Pearson’s correlation coefficient were then computed to assess the relationship between early activity and performance.

## Supplementary Figures

**Fig. S1.**
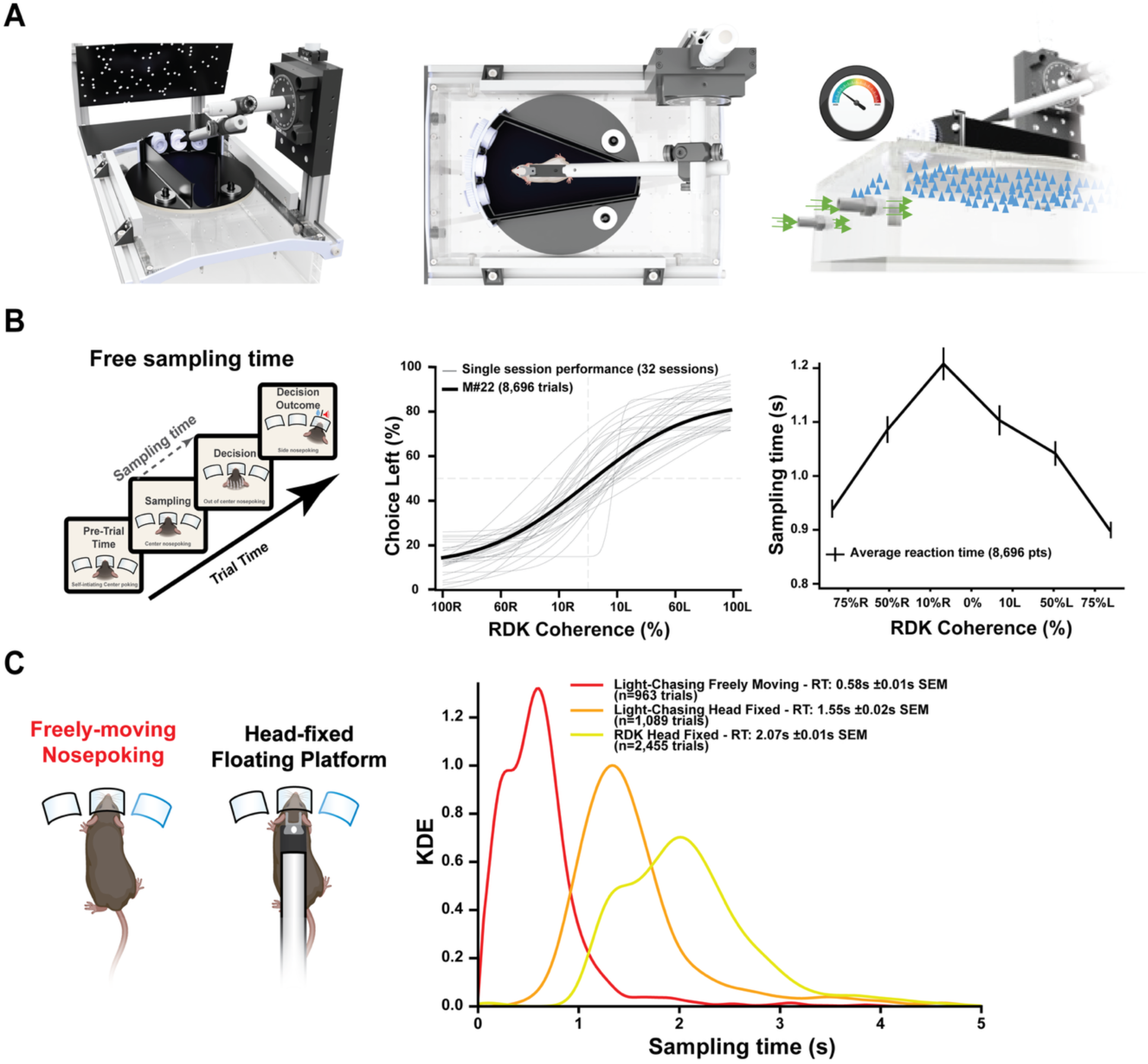
A novel floating-platform nose-poke system to study neural mechanisms of decision-making in head-fixed mice. (**A**) 3D illustrations showing side (left), top (middle), and schematic bottom (right) views of the nose-poke floating-platform setup designed for head-fixed mice performing a Random Dots Kinematogram (RDK) task. (**B**) Free-sampling RDK task structure (left), psychometric curves showing choice performance (% choice-left) across individual sessions for an example mouse (middle; thin gray lines: single sessions, bold black line: average), and average sampling time as a function of stimulus coherence levels for rightward (R) and leftward (L) choices (right). (**C**) Left: Schematics illustrating experimental setups comparing freely-moving (nose-poking) and head-fixed (nose-poking floating-platform) paradigms. Right: Kernel density estimates (KDE) of sampling-time distributions comparing an example mouse performing freely-moving light-chasing (red), head-fixed light-chasing (orange), and head-fixed RDK (yellow) tasks. Mean reaction times ± SEM indicated.

**Fig. S2.**
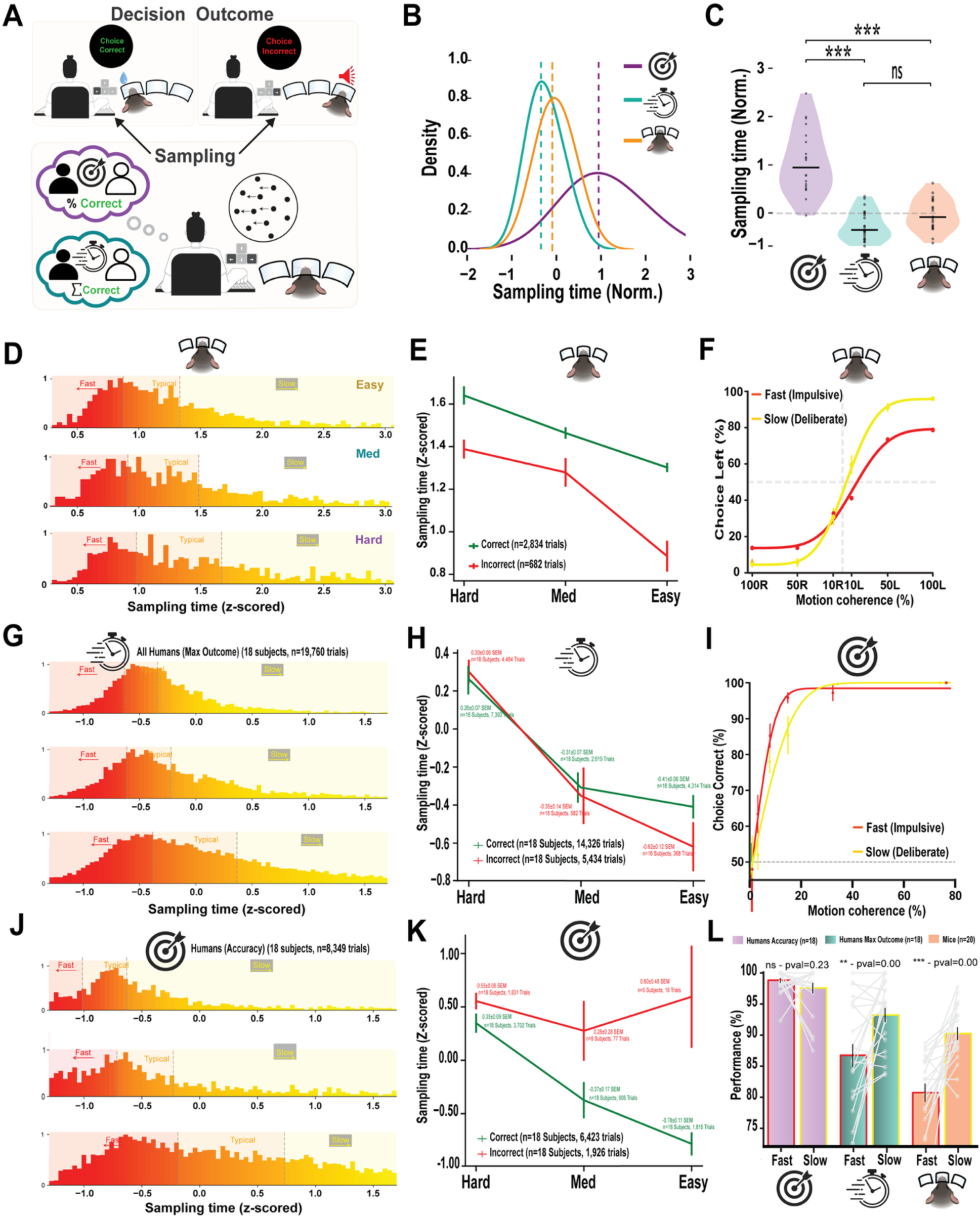
Switching between fast and slow behavior is a common strategy during voluntary behavior in both humans and mice. (**A**) Schematic of the human RDK task. Participants performed the task in two contexts: Context I (Accuracy)—emphasizing correct performance, and Context II (Speed)—emphasizing maximizing correct responses. Subjects responses, reported by pressing left or right arrow keys, were followed by outcome feedback. (**B**) Distribution of normalized sampling times (z-scored) across contexts. Violet: human accuracy context; turquoise: human speed context; orange: head-fixed mice on the floating platform. Dashed lines indicate means for each distribution. (**C**) Sampling times were significantly longer in the human accuracy context compared to both the human speed and mouse conditions. Sampling time distributions in human speed and mouse contexts were not significantly different (ns). (**D**) Sampling time histograms (seconds) across RDK stimulus difficulties (easy, medium, hard) for an example mouse (M#85, 3,519 trials), subdivided into fast, typical, and slow trials. (**E**) Mean sampling time (seconds) for correct (green) and incorrect (red) trials across difficulty levels (hard, medium, easy) for the same example mouse. (**F**) Psychometric performance (choice-left %) as a function of motion coherence for fast (red) and slow (yellow) trials. (**G–H**) Data from human participants in the speed context (n=22). (**G**) Sampling time distributions (z-scored) across difficulties for fast, typical, and slow trials. **H**) Mean sampling time (z-scored) for correct (green) vs. incorrect (red) trials across difficulty levels. (**I**) Psychometric curve showing choice accuracy across motion coherences for fast (red) vs. slow (yellow) trials during human accuracy context (n = 20). (**J–K**) Same as panels G–H, but for human participants in the accuracy context (n = 20). (**J**) Sampling time histograms by difficulty level. (**K**) Sampling time comparison between correct and incorrect trials across difficulty. (**L**) Accuracy during easy trials for fast (red) and slow (yellow) decisions across three groups: human accuracy context, human speed context, and head-fixed mice. Grey lines represent individual subjects or mice; bars indicate group means.

**Fig. S3.**
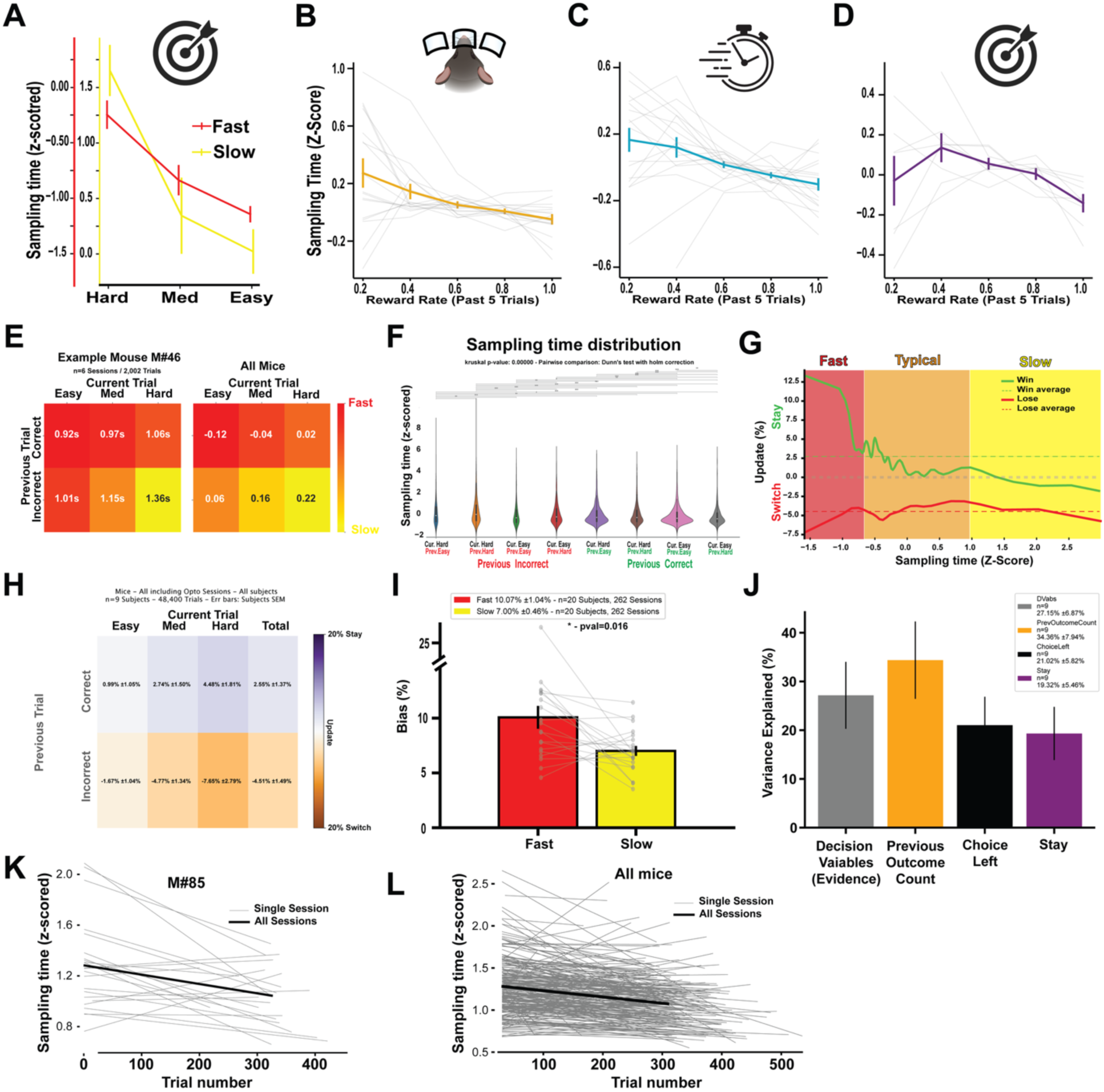
Trial history and decision outcome and sensory evidence modulate sampling time. (**A**) Mean sampling time (z-scored) across stimulus difficulties during fast (red) and slow (yellow) behavioral strategies humans under the accuracy context. (**B–D**) Sampling time (z-scored) as a function of recent reward rate (averaged over the preceding 5 trials) in: (**B**) Head-fixed mice (orange), (**C**) Human subjects under the speed context (turquoise), and (**D**) Human subjects under the accuracy context (violet). Thin gray lines represent individual animals or participants; bold lines show group means. (**E**) Heatmap showing average sampling time as a function of previous trial outcome (correct/incorrect) and current trial difficulty. Left: example mouse (M#46); Right: mean across all mice. Warmer colors indicate longer sampling times. (**F**) Violin plots illustrating distributions of sampling time (z-scored) across combinations of previous trial outcome (correct/incorrect) and current trial difficulty. Trial history and trial difficulty significantly affect sampling time. (**G**) Probability of win (green) and lose (red) behavioral updates as a function of sampling time (z-scored). Fast decisions are more likely to be driven by outcome-insensitive strategies. (**H**) Impact of previous trial outcome (correct/incorrect) on win /lose behavior update across current trial difficulties (easy, medium, hard). Percentages indicate update probabilities. (**I**) Choice bias (preference for one direction) is higher during fast strategies compared to slow strategies across mice. Paired comparisons across animals reveal significant reduction in bias during slow decisions (*p* = 0.016). (**J**) Variance in sampling time explained (OLS model) by different task variables, including decision evidence (stimulus strength), previous outcome, previous choice, and stay/switch behavior. (**K**) Linear regression of sampling time (seconds) over trials for an example mouse (M#85), showing reduced sampling over the course of a session. Each gray line represents a single session; the bold line shows the fit across sessions. (**L**) Same as panel K, averaged across all mice.

**Fig. S4.**
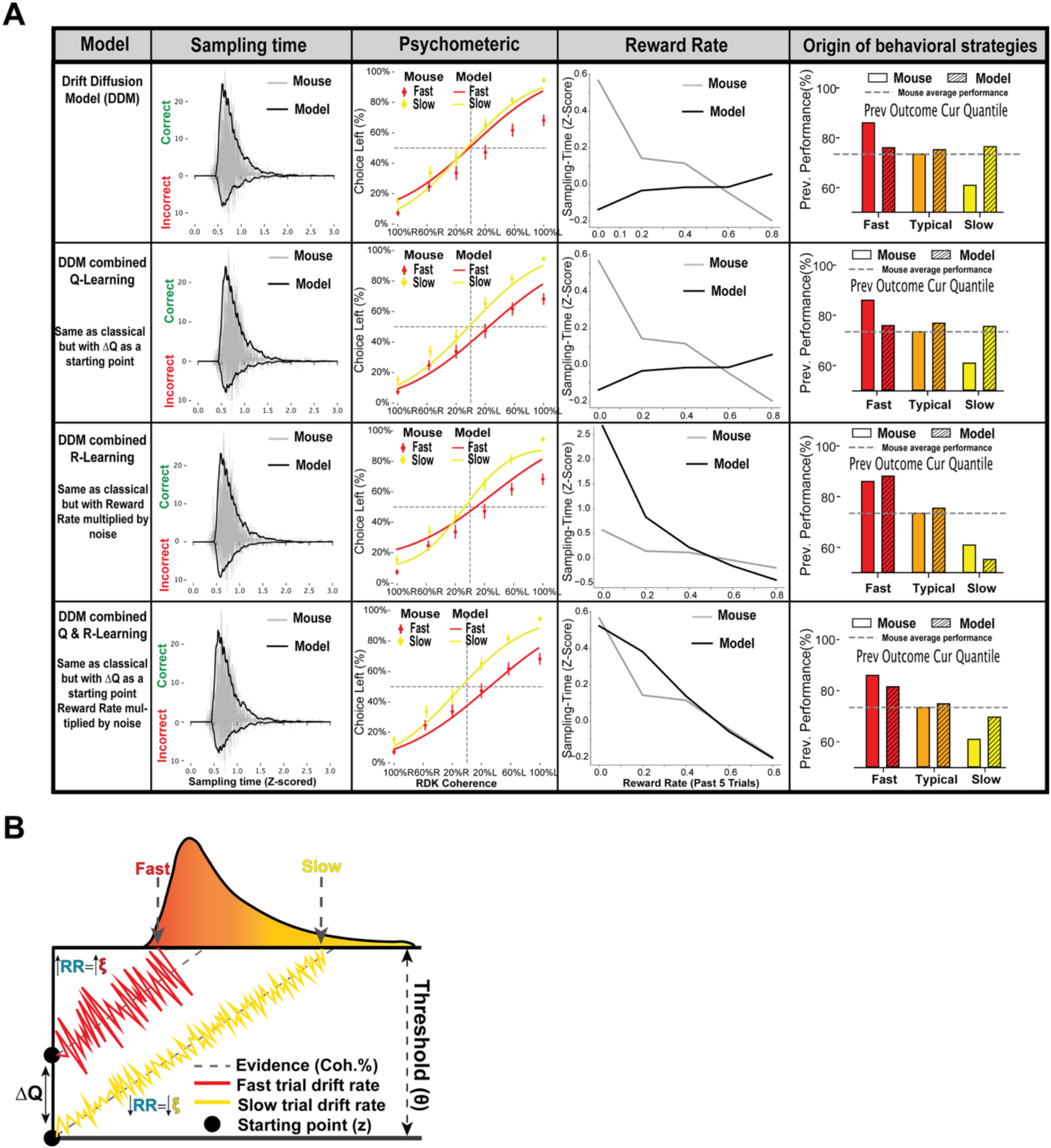
Reinforcement learning (RL)-combined drift diffusion models (DDMs) capture distinct behavioral strategies during fast and slow decisions. (**A**) Four computational models were fit to mouse correct and incorrect trial distribution, each panel (left to right) reflecting sampling time distributions, psychometric performance, reward rate sensitivity, and trial-history dependencies: (Top to bottom) 1. Classic Drift Diffusion Model (DDM). 2. Q-DDM: DDM with ΔQ-learning action values incorporated as a starting point bias. 3. R-DDM: DDM with R-learning value (reward rate) modulating noise to affect drift rate. 4. Q&R-DDM: Combined model with ΔQ-learning action values setting the starting point and R-learning value modulating the drift rate. Each model (black) is fit to mouse data (grey) and the resulting distributions are shown: Sampling time distributions (seconds) for correct and incorrect trials, psychometric performance (choice left%) across motion coherence levels during fast (red) and slow (yellow) trials, reward rate as a function of sampling time, and previous trial outcome vs. current behavioral strategy (fast, typical, slow), showing model vs. mouse performance dependence on history. (**B**) Schematic illustrating how RL variables are integrated into a classical DDM framework to explain behavioral strategy. Evidence (coherence %) is accumulated toward a decision threshold (θ). ΔQ-learning action values biases the starting point (z), affecting early decision tendencies. R-learning value modulates noise amplitude, influencing the drift rate (slope), with lower reward rates yielding more stable accumulation in slow trials and more variable trajectories in fast trials.

**Fig S5.**
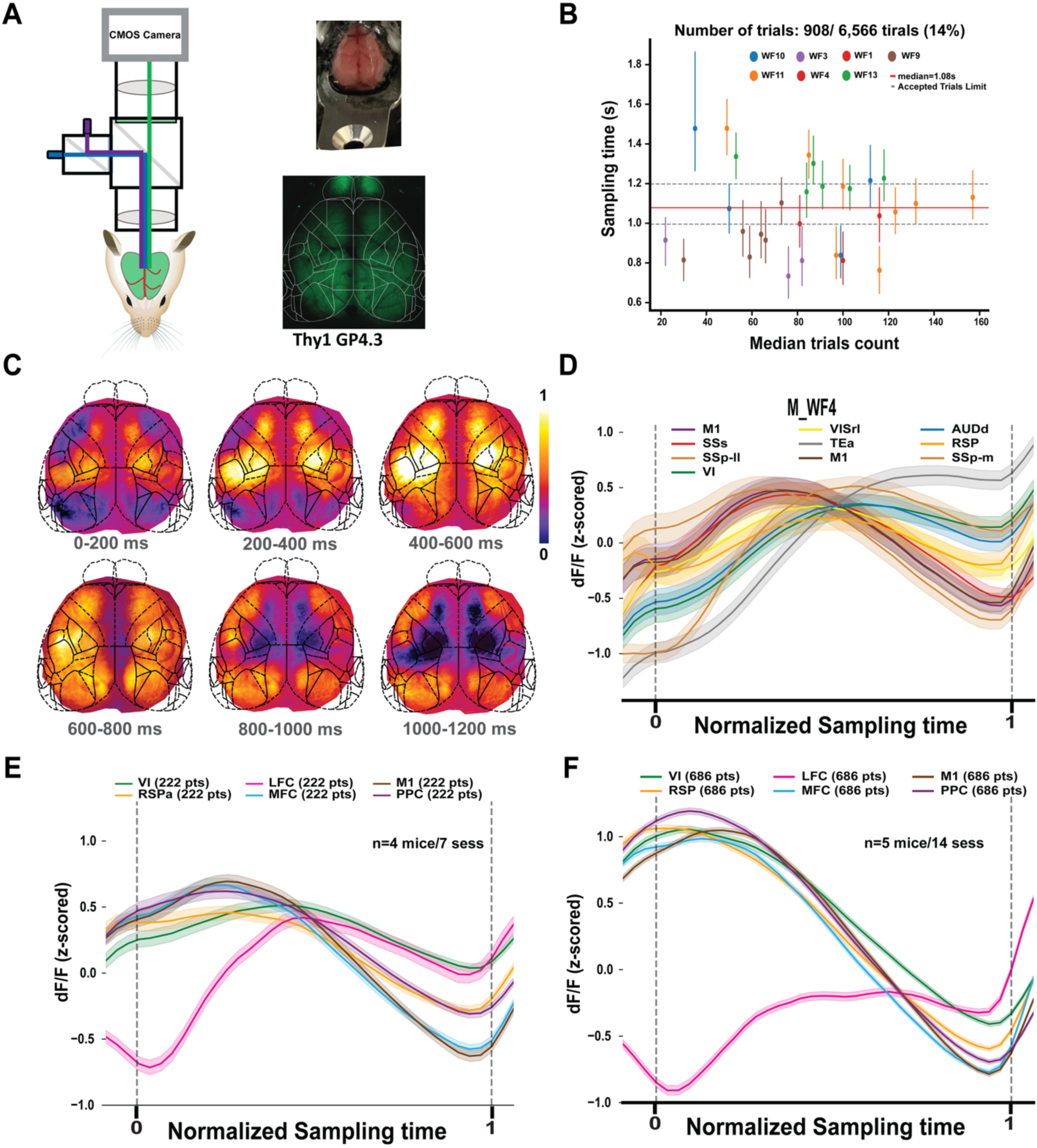
Widefield calcium imaging reveals cortical dynamics during typical trials across mice. **(A)** Schematic of the widefield imaging setup for monitoring neural activity across the dorsal cortex (left). Top right: clear skull under white light illumination. Bottom right: same view under blue light for calcium imaging in Thy1-GCaMP-expressing mice. **(B)** Overview of selected imaging sessions across mice focused on typical trials with sampling durations between 1000–1200 ms. Each point indicates a session, colored by mouse identity; dashed lines indicate inclusion thresholds. **(C)** Pixel-wise heatmaps from an example mouse show spatiotemporal dynamics of calcium activity during the sampling period in typical-duration trials, averaged over successive 200-ms windows. **(D)** Trial-averaged calcium dynamics from selected cortical areas in the example mouse, aligned to normalized sampling time. **(E)** Cross mice average of calcium dynamics from multiple cortical regions (n=4 mice, 7 sessions), showing consistent spatiotemporal frontal dynamics frontal during typical decisions. **(F)** Cross mice (n=5 mice, 14 sessions), exhibit similar spatiotemporal frontal dynamics similar (E). However, most cortical areas exhibit a noticeable decay in activity before the end of sampling.

**Fig. S6.**
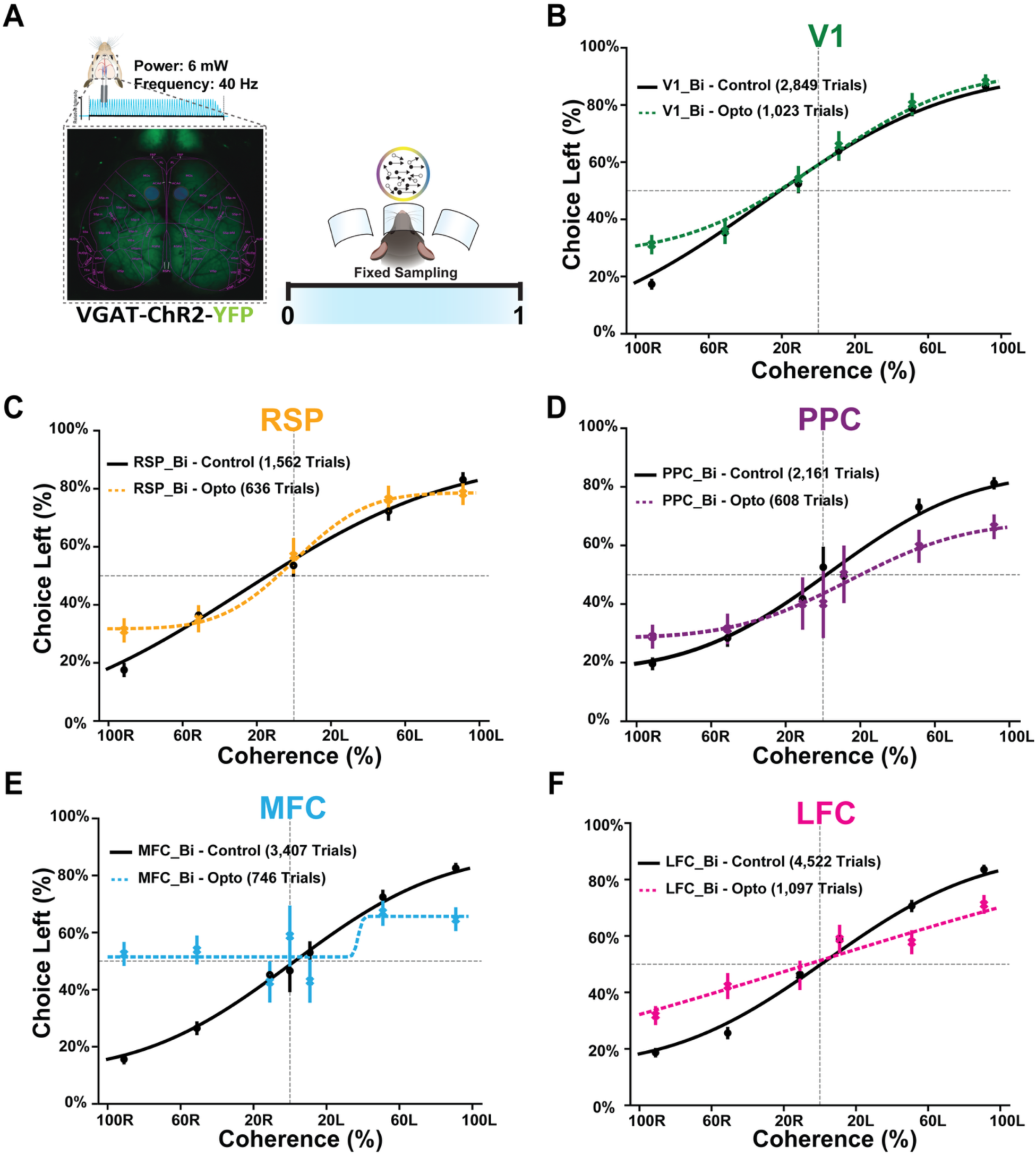
Optogenetic inactivation of distinct cortical areas during fixed sampling paradigm reveals region-specific contributions to decision-making. (**A**) Schematic showing bilateral optogenetic inactivation during a fixed 1-second sampling window in VGAT-ChR2-YFP mice. Inactivation was delivered at 6 mW and 40 Hz using chronically implanted optical fibers targeting specific cortical areas under a clear skull preparation. (**B–F**) Psychometric performance curves comparing control trials (solid black lines) and optogenetic inactivation trials (dashed lines) across motion coherence levels for each targeted brain region: (**B**) Primary visual cortex (V1, green), (**C**) Retrosplenial cortex (RSP, orange), (**D**) Posterior parietal cortex (PPC, violet), (**E**) Medial frontal cortex (MFC, blue), (**F**) Lateral frontal cortex (LFC, magenta). Data show a region-specific effect of inactivation on perceptual performance, with frontal areas (particularly MFC and LFC) showing more pronounced impairments of behavior compared to early sensory areas.

**Fig. S7.**
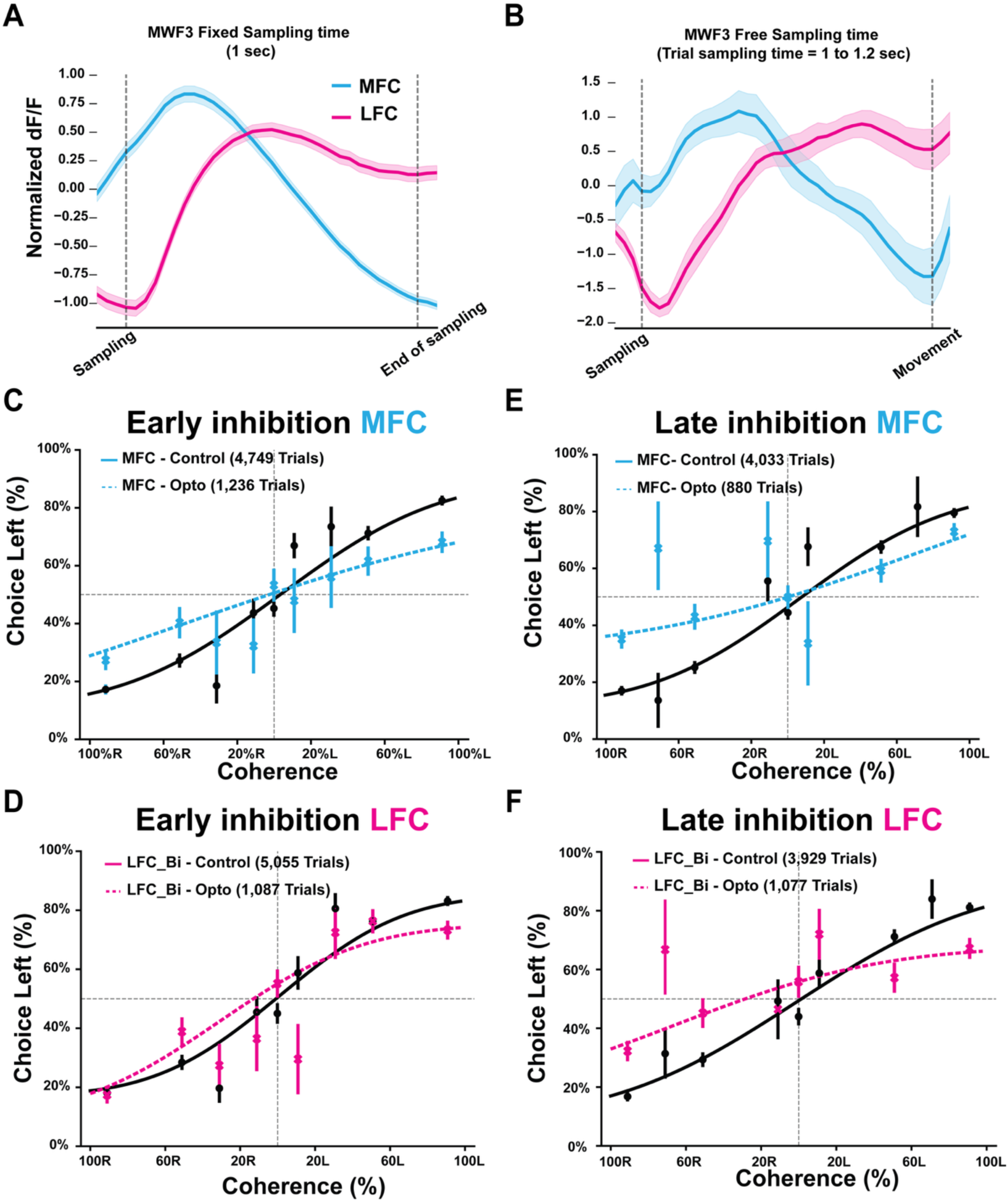
Frontal cortex spatiotemporal dynamics and its causal contributions to decision-making during sampling. (**A**) Frontal cortical calcium dynamics during a fixed 1-second sampling task, showing normalized widefield fluorescence from medial frontal cortex (MFC, blue) and lateral frontal cortex (LFC, magenta) in an example mouse. MFC shows an earlier, more transient peak, while LFC exhibits delayed and sustained activity. (B) Same as (A), but during free sampling trials with durations between 1 and 1.2 seconds for the same mouse. The spatiotemporal activity pattern across MFC and LFC is preserved and reflects voluntary sampling dynamics. (**C–F**) Psychometric performance across different difficulties during optogenetic inhibition at early (0–350 ms) and late (650– 1000 ms) phases of the sampling period. (**C, D**) Early inhibition in MFC (C, blue) and LFC (D, magenta). (**E, F**) Late inhibition in MFC (E) and LFC (F). Solid lines: control trials; dashed lines: optogenetic inhibition trials. Data reveal timing- and region-specific effects on perceptual decisions, with MFC inhibition having the most pronounced impact across all time.

**Fig. S8.**
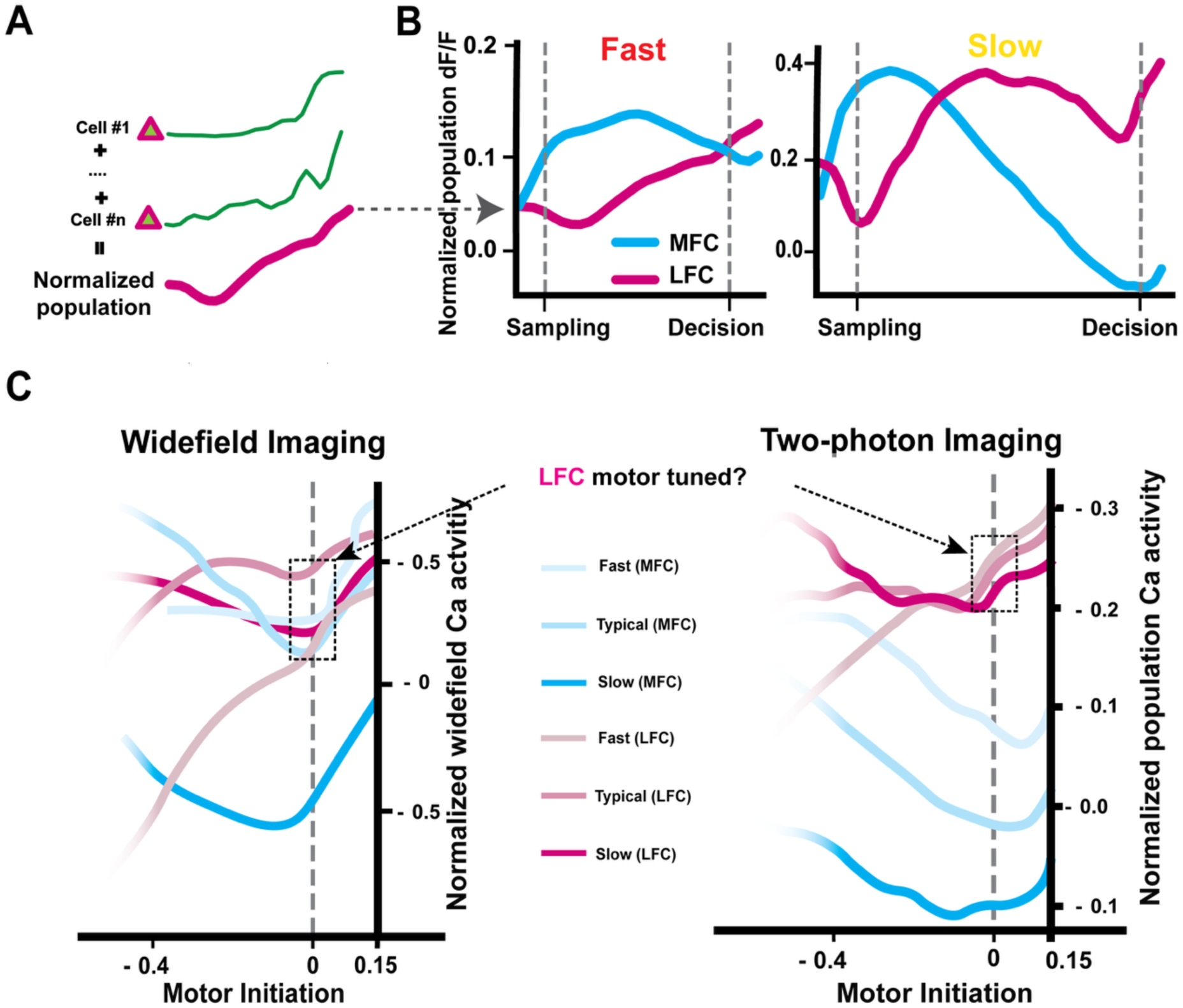
Consistent spatiotemporal calcium dynamics in layer 2/3 frontal neurons across behavioral strategies. (**A**) Schematic illustrating calculating population-averaged calcium responses from individual neurons. Left: trial-aligned calcium traces from example L2/3 neurons. Right: resulting normalized population activity over time. (**B**) Normalized population calcium responses of L2/3 population from medial frontal cortex (MFC, blue) and lateral frontal cortex (LFC, magenta) during fast (left) and slow (right) trials in an example mouse. While MFC activity peaks early in fast trials, LFC shows delayed and more sustained activity, particularly during slow trials. (**C**) Comparison of frontal cortex calcium dynamics aligned to motor initiation, using both widefield imaging (left) and two-photon imaging (right). In both methods, LFC shows a ramping activity pattern consistent with motor-related tuning, especially in slow and typical trials. MFC activity peaks earlier and declines before movement onset.

**Fig. S9.**
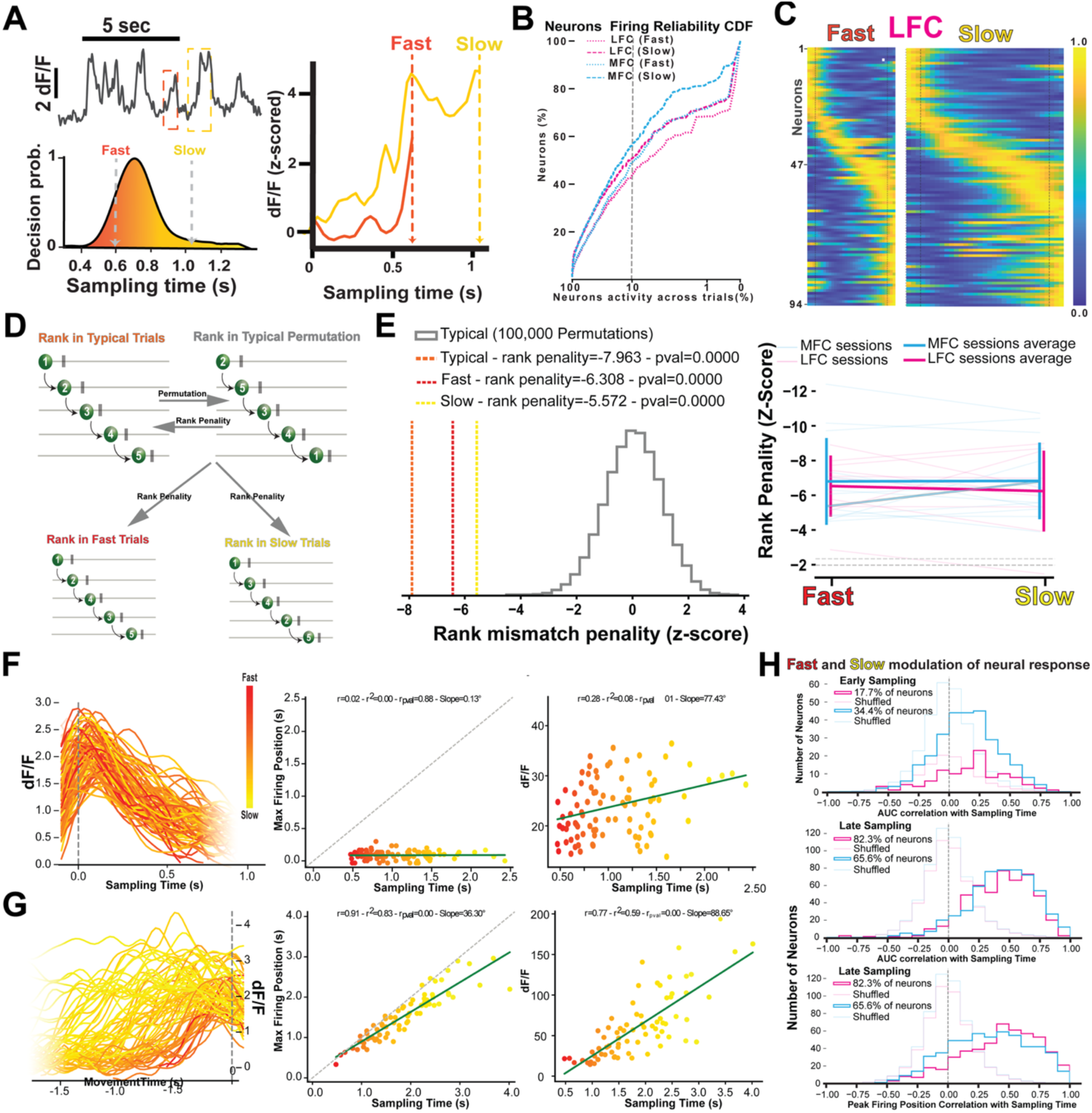
Temporal organization of L2/3 frontal neuron sequences during fast and slow behavioral strategies. (**A**) Top: Single-neuron calcium traces showing fast and slow decision epochs. Bottom left: Decision probability distribution for fast (red) and slow (yellow) trials in an example session. Bottom right: Average z-scored dF/F trace for fast and slow trials aligned to sampling time. (**B**) Cumulative distribution function (CDF) of trial-by-trial firing reliability for L2/3 neurons in MFC and LFC during fast and slow trials. (C) Heatmap of calcium activity from LFC neurons in an example session, aligned by average peak response across trials during fast (left) and slow (right) strategies. Bottom: Rank penalty scores across all MFC and LFC sessions showing consistent differences between strategies. (**D**) Schematic of the permutation-based approach for computing rank mismatch penalties across fast, typical, and slow trials. Neuronal sequences from each strategy are compared to permuted typical-trial sequences as a reference. (**E**) Left: Histogram showing rank mismatch penalties for typical (orange), fast (red), and slow (yellow) trials compared to permuted typical-trial sequences from a single example session. Right: Summary across all sessions showing consistent sequence deviation in all three strategies (see methods). (**F**) Example L2/3 neuron responses across trials (color-coded from red (fast) to yellow (slow) by sampling duration), aligned to sampling onset. Left: single trials dF/F traces. Middle: Correlation between sampling duration and peak time. Right: Correlation between sampling duration and response amplitude (AUC). (**G**) Same example neuron as in F, but responses aligned to movement onset. Left: single trials dF/F traces. Middle: Sampling duration vs. peak time. Right: Sampling duration vs. AUC. Both timing and amplitude are significantly modulated by sampling duration. (**H**) Population-level analysis showing distributions of correlation coefficients between sampling time and neuronal responses: Top: AUC during early sampling (<0.2 seconds of sampling time). Middle: AUC during late sampling (>0.2 seconds of sampling time). Bottom: Peak response timing during late sampling. Blue: MFC neurons; magenta: LFC neurons; grayed blue and magenta are shuffled controls.

**Fig. S10.**
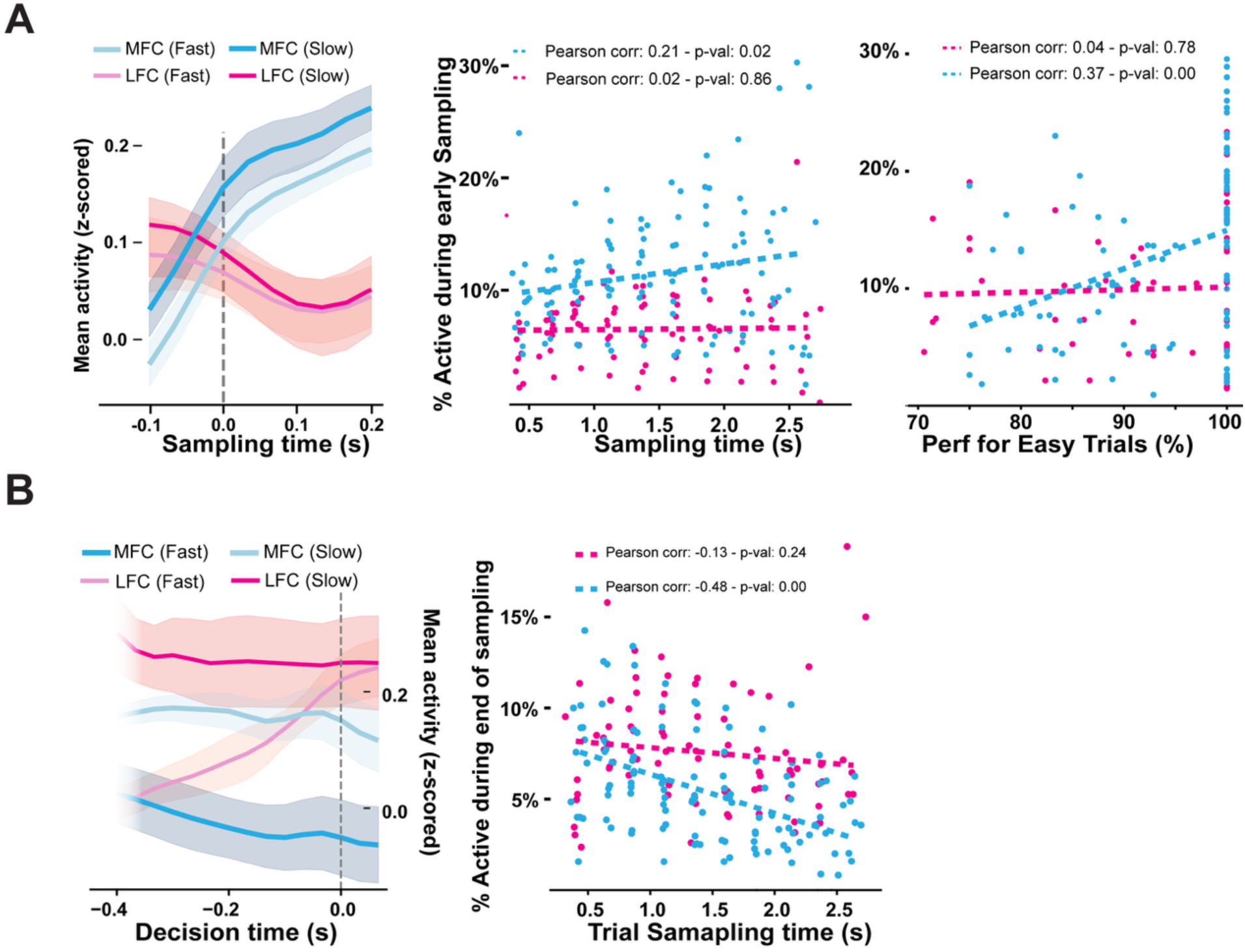
Frontal L2/3 population activity is differentially tuned to sampling time and performance in MFC but not LFC. **A)** Left: Mean z-scored population activity in MFC (blue) and LFC (magenta) aligned to the onset of sampling, separated by fast (faint lines) and slow (dark lines) behavioral strategies. Middle: Percentage of active L2/3 neurons during the early phase of sampling as a function of sampling time across mice and sessions. Dashed lines show linear fits with Pearson correlation coefficients and p-values. Right: Correlation between early sampling activity and behavioral performance (mean % correct) during easy trials across mice and sessions (see methods). **B**) Left: Mean z-scored population activity in MFC and LFC across mice and sessions aligned to the end of sampling (just before movement initiation), separated by fast and slow trials. Right: Percentage of neurons active at the end of sampling as a function of sampling time across mice and sessions (see methods). MFC activity shows a stronger negative correlation with trial duration than LFC, indicating tighter tuning to behavioral timing in MFC.

**Fig. S11.**
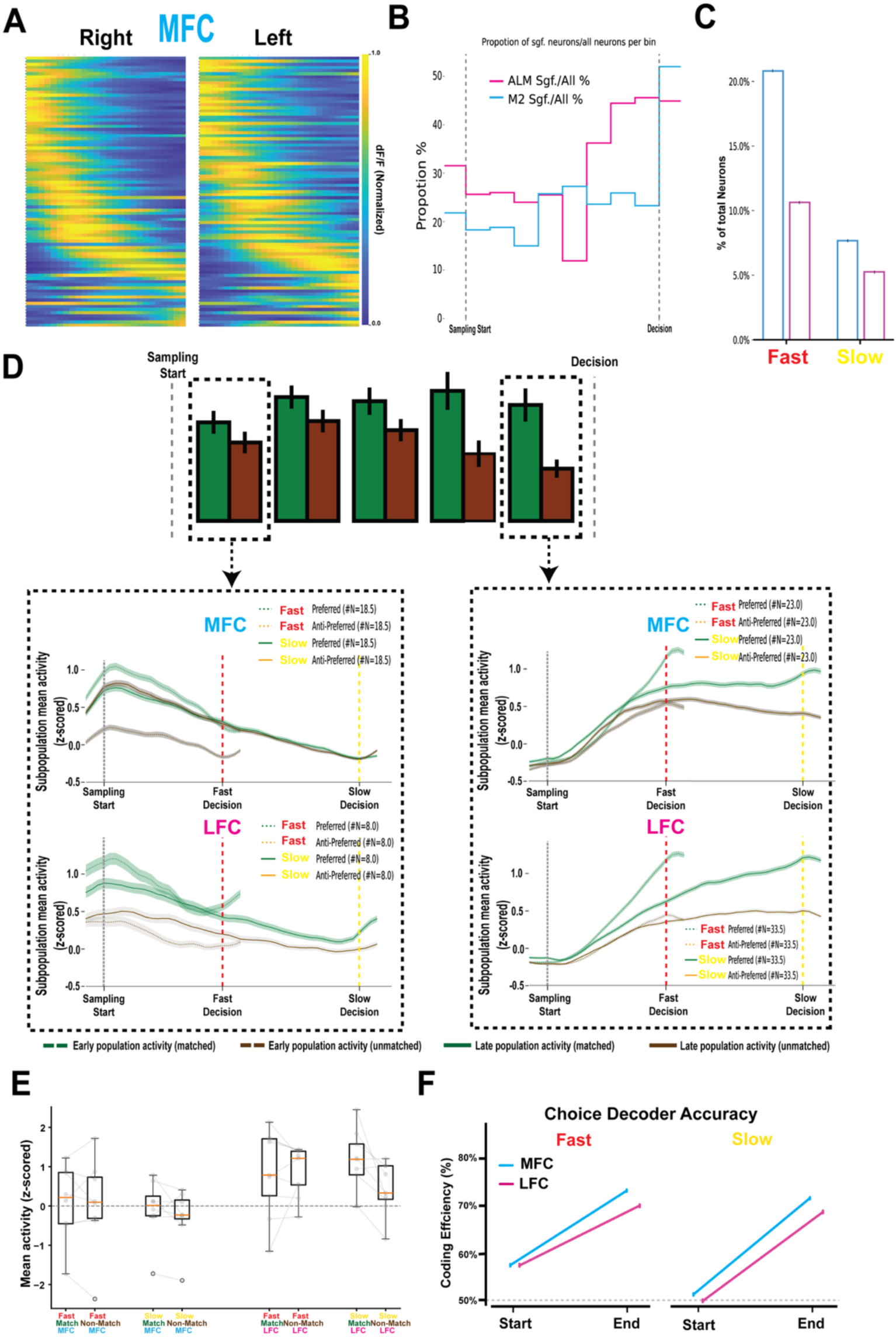
Choice-selective neural population dynamics during fast and slow decisions. (**A**) Heatmaps of normalized calcium activity from MFC neurons in an example session for rightward choices (left panel) and leftward choices (right panel), illustrating population sequence alignment is consistent for left and right choices. (**B**) Proportion of choice-selective neurons over normalized sampling time in MFC (blue) and LFC (magenta). LFC neurons show delayed encoding compared to MFC, which encode choice homogeneously across sampling time. (**C**) Percent of choice encoding neurons at the start of sampling during fast and slow trials (see methods). (**D**) Top: Population activity of choice-preferring (green) and non-preferring (brown) neural subgroups at the start (dashed box) and end (dashed box) of sampling. Bottom: Subpopulation mean activity for matched and unmatched subgroups in MFC (top) and LFC (bottom), showing divergence in dynamics between fast and slow strategies at the start and end of sampling. (**E**) Difference in mean population activity (z-scored) between the last neural bin before the decision and the first neural bin at the onset of sampling. Values are shown separately for MFC and LFC, and across matched vs. non-matched choice-neurons during fast and slow trials. This metric captures the net buildup or declines in activity from initial sensory sampling to the moment of choice commitment. (**F)** Choice decoder accuracy (coding efficiency) computed at the start and end of sampling, separately for fast and slow trials. Both MFC (blue) and LFC (magenta) show improved decoding performance toward the end of the sampling time.

**Fig. S12.**
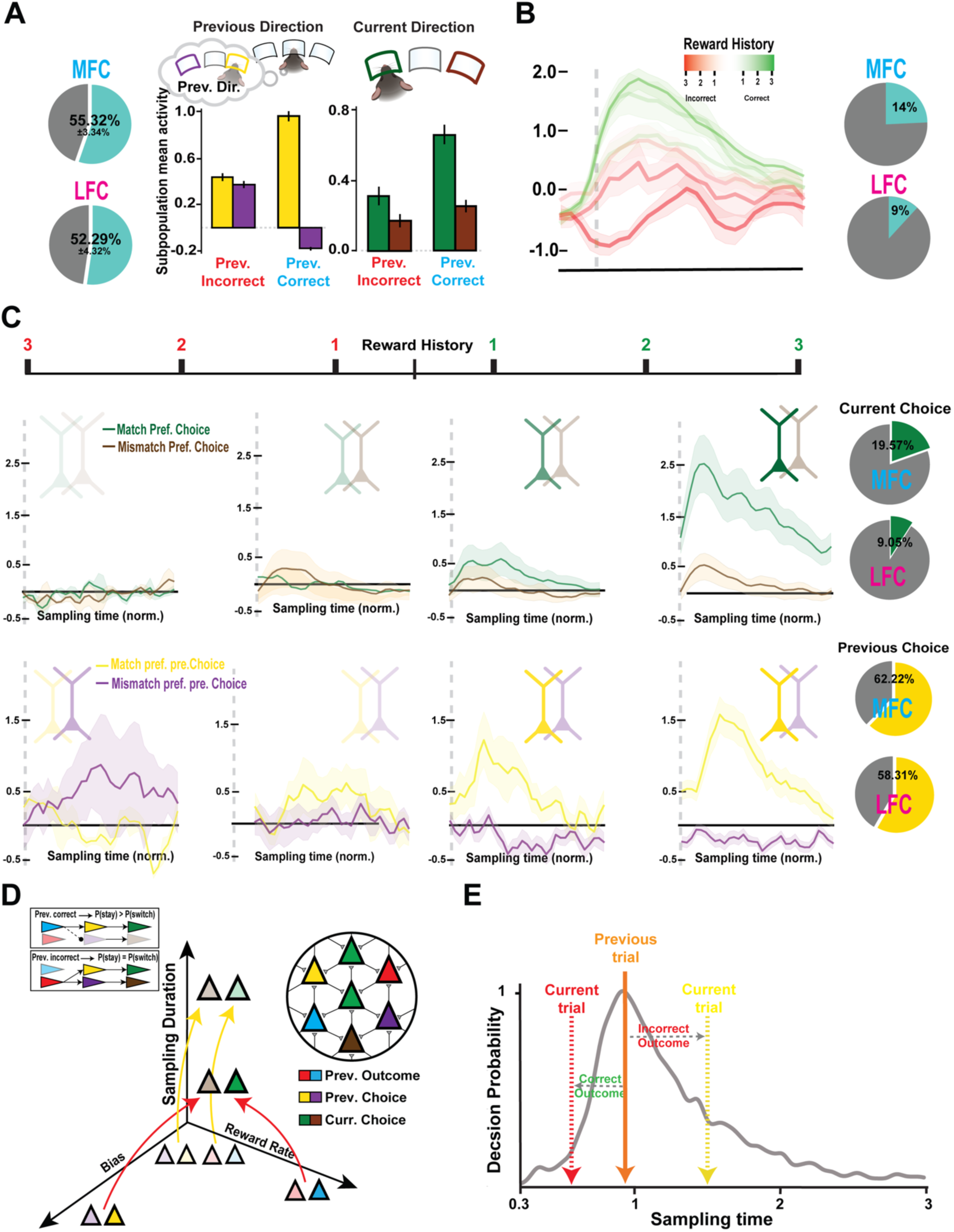
Previous trial outcomes and reward history modulate frontal cortex encoding of current and past choices. (**A**) **Left:** Proportion of outcome-sensitive neurons in MFC and LFC. Right: The influence of previous outcome on neurons selective for previous choice direction (left) and current choice direction (right) at the start of sampling. (**B**) Left: Example neuron activity is modulated by recent reward history (reward rate over the last 3 trials), aligned to normalized sampling time. Right: Pie charts showing the proportion of reward-rate-sensitive neurons in MFC and LFC. (**C**) Top: Example neurons encoding the current choice, showing modulation by recent reward rate. Bottom: Example neurons encoding the previous choice, also modulated by recent reward rate. Right: Pie charts showing the proportion of current choice (green) and previous choice (yellow) encoding neurons modulated by previous outcome. (**D**) Schematic illustrating how outcome and choice history jointly influence decision strategy. (**E**) Schematic representation of how sampling time variability reflects history-dependent strategy shifting.

**Fig. S13.**
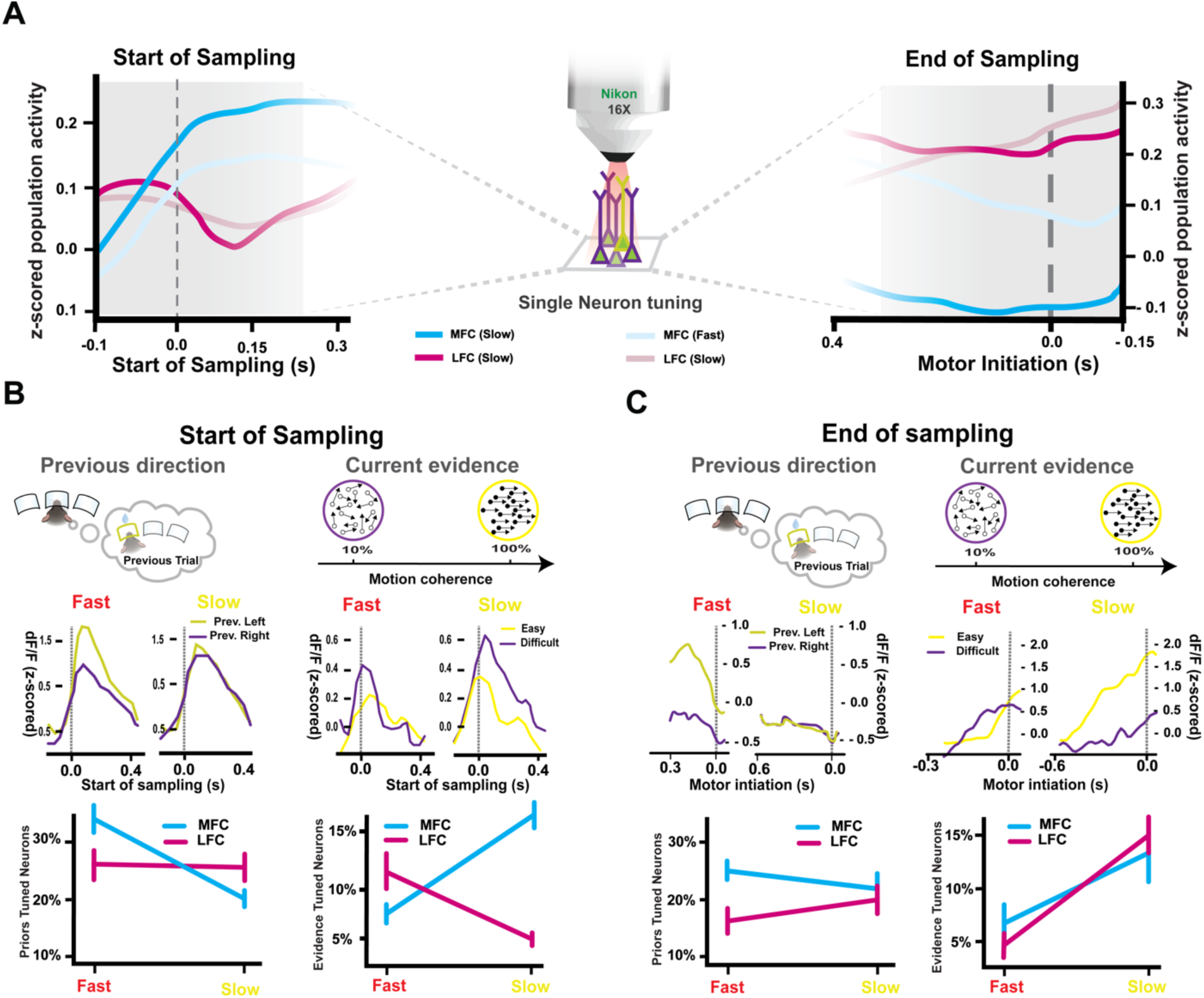
Neural representation of trial history and evidence during fast and slow behavior. **(A**) Population activity (z-scored) in MFC (blue) and LFC (magenta) during fast (impulsive, light lines) and slow (deliberate, dark lines) trials. Left: Aligned to the start of sampling, showing stronger early activation in MFC during slow trials. Right: Aligned to motor initiation (end of sampling), showing greater LFC activation prior to movement. Center: schematic of the two-photon imaging setup used for single-neuron resolution. (**B**) Neural tuning at the start of sampling for two decision-related variables: Left column: Previous trial’s choice direction (left vs. right). Right column: Current trial’s stimulus strength (easy vs. difficult motion coherence). Top: Example neuron responses for fast and slow strategies. Bottom: Percentage of significantly tuned neurons in MFC and LFC during fast and slow (see methods). (**C**) Same analysis as panel B, but for the end of sampling, aligned to movement initiation.

**Fig. S14.**
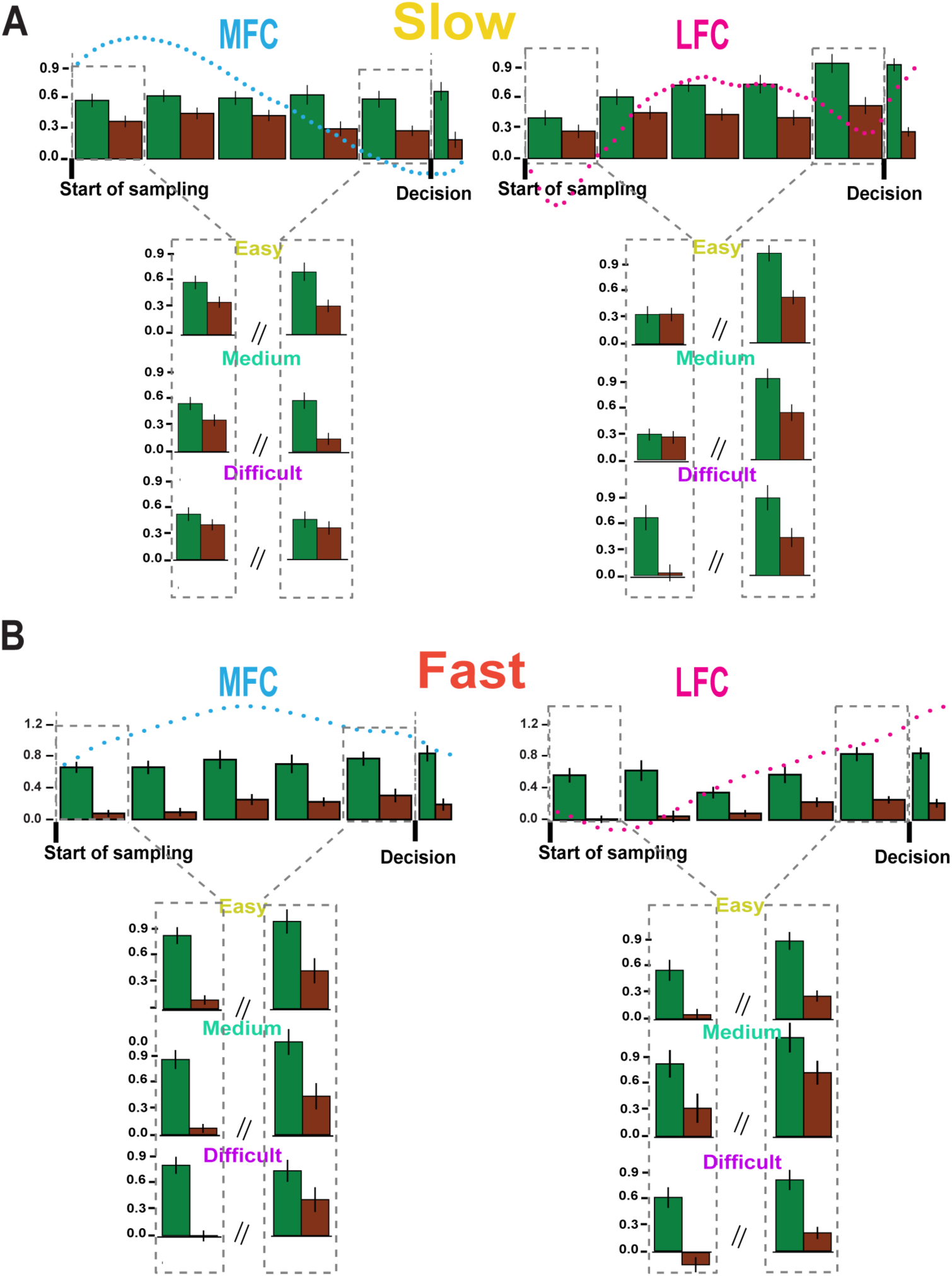
Choice coding efficiency reflects sensitivity to sensory evidence across behavioral strategies. (**A**) **Top:** Sequence-aligned choice-selective neuron responses in MFC (left) and LFC (right) during slow trials, plotted as mean population activity for matched-choice (green) vs mismatch (brown) conditions, aligned to different temporal bins along sampling time. Bottom: Breakdown of neural activity by stimulus difficulty (easy, medium, difficult) for both matched and mismatch trials at the start and end of sampling. During slow decisions, matched-choice neurons show stronger evidence-related tuning, particularly in MFC. (**B**) Same analysis as in (**A**), but for fast trials. Matched-choice neuron activity is more similar across difficulties..

**Fig. S15.**
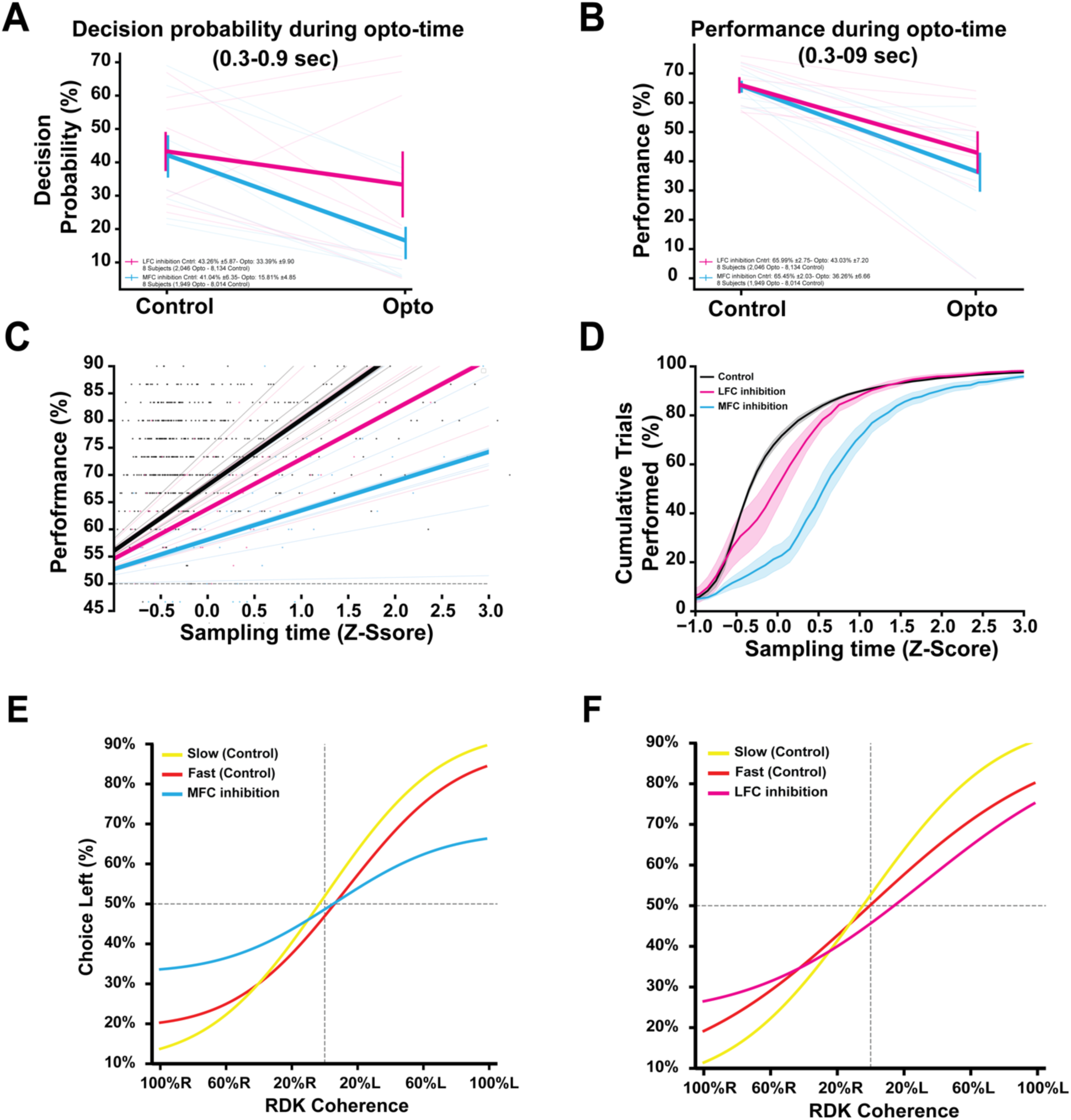
Mid-sampling inhibition of MFC and LFC disrupts decision probability, accuracy during free sampling paradigm. (**A**) Decision probability during the optogenetic inhibition window (0.3–0.9 s after sampling onset) significantly decreased with **MFC** (blue) and **LFC** (magenta) inhibition. (**B**) Task performance during optogenetic inhibition showed a significant drop under both MFC and LFC inhibition compared to control. (**C**) Linear fit of task performance as a function of z-scored sampling time, separated by condition: control (black), MFC inhibition (blue), and LFC inhibition (magenta). MFC inhibition flattens the performance-sampling time relationship. (**D**) Cumulative trial distribution across z-scored sampling time. MFC inhibition delays the accumulation of decisions compared to control and LFC inhibition. (**E–F**) Psychometric curves show behavioral performance across RDK motion coherence in control slow (yellow), control fast (red), and MFC-inhibited (E, blue) or LFC-inhibited (F, magenta) trials.

---

**Video S1. Head-fixed mice navigate a nose-poking floating platform.** Side view of a head-fixed mouse controlling a circular floating platform equipped with three nose-pokes. Each trial begins when the mouse initiates a nose-poke in the center port, triggering the presentation of a random-dot kinematogram (RDK) stimulus. The mouse then voluntarily withdraws from the center port to terminate stimulus sampling and rotates the platform to reach either the left or right choice port to report its decision and collect a reward.

**Video S2. Head-fixed mice display both fast and slow sampling strategies under identical sensory conditions.** Top-view of a head-fixed mouse performing on a nose-poking platform during a decision-making task. The video shows two representative trials at intermediate stimulus difficulty (50% coherence), illustrating that mice can adopt either fast or slow sampling strategies for the same sensory evidence.

**Video S3. Head-fixed mice exhibit variable sampling durations while maintaining robust performance during RDK task in nose-poking floating platform.** Top left: Psychometric function from an example behavioral session, showing performance across stimulus difficulties. Top right: Histogram of sampling time distribution from the same session, illustrating within-session variability in sampling behavior. Bottom: Top-down video of a head-fixed mouse engaged in the decision-making task on the nose-poking platform.

## References

1. D. Kahneman, Thinking, Fast and Slow (Farrar, Straus and Giroux, New York, NY, US, 2011)Thinking, fast and slow.

2. D. Luce, “Response Times: Their Role in Inferring Elementary Mental Organization” (1986; https://www.semanticscholar.org/paper/Response-Times%3A-Their-Role-in-Inferring-Elementary-Luce/4f714b26b7ceb23ccad0e47d19a6fd94ec10bf4f).

3. R. Bogacz, E. Brown, J. Moehlis, P. Holmes, J. D. Cohen, The physics of optimal decision making: A formal analysis of models of performance in two-alternative forced-choice tasks. Psychological Review 113, 700–765 (2006).

4. R. Ratcliff, G. McKoon, The Diffusion Decision Model: Theory and Data for Two-Choice Decision Tasks. Neural Comput 20, 873–922 (2008).

5. J. I. Gold, M. N. Shadlen, The Neural Basis of Decision Making. Annual Review of Neuroscience 30, 535–574 (2007).

6. P. W. Glimcher, The Neurobiology of Visual-Saccadic Decision Making. Annual Review of Neuroscience 26, 133–179 (2003).

7. M. A. Nashaat, H. Oraby, R. N. S. Sachdev, Y. Winter, M. E. Larkum, Air-Track: a real-world floating environment for active sensing in head-fixed mice. Journal of Neurophysiology 116, 1542–1553 (2016).

8. S. E. Dominiak, M. A. Nashaat, K. Sehara, H. Oraby, M. E. Larkum, R. N. S. Sachdev, Whisking Asymmetry Signals Motor Preparation and the Behavioral State of Mice. J. Neurosci. 39, 9818–9830 (2019).

9. J. Voigts, M. T. Harnett, Somatic and Dendritic Encoding of Spatial Variables in Retrosplenial Cortex Differs during 2D Navigation. Neuron 105, 237–245.e4 (2020).

10. T. Marques, M. T. Summers, G. Fioreze, M. Fridman, R. F. Dias, M. B. Feller, L. Petreanu, A Role for Mouse Primary Visual Cortex in Motion Perception. Current Biology 28, 1703–1713.e6 (2018).

11. M. N. Shadlen, R. Kiani, Decision Making as a Window on Cognition. Neuron 80, 791–806 (2013).

12. L. Hernández-Navarro, A. Hermoso-Mendizabal, D. Duque, J. de la Rocha, A. Hyafil, Proactive and reactive accumulation-to-bound processes compete during perceptual decisions. Nat Commun 12, 7148 (2021).

13. C. Schwarz, H. Hentschke, S. Butovas, F. Haiss, M. C. Stüttgen, T. V. Gerdjikov, C. G. Bergner, C. Waiblinger, The head-fixed behaving rat—Procedures and pitfalls. Somatosensory & Motor Research 27, 131–148 (2010).

14. J. I. Sanders, A. Kepecs, Choice ball: a response interface for two-choice psychometric discrimination in head-fixed mice. Journal of Neurophysiology 108, 3416–3423 (2012).

15. B. a. J. Reddi, R. H. S. Carpenter, The influence of urgency on decision time. Nat Neurosci 3, 827–830 (2000).

16. J. Palmer, A. C. Huk, M. N. Shadlen, The effect of stimulus strength on the speed and accuracy of a perceptual decision. Journal of Vision 5, 1 (2005).

17. J. Drugowitsch, R. Moreno-Bote, A. K. Churchland, M. N. Shadlen, A. Pouget, The Cost of Accumulating Evidence in Perceptual Decision Making. J. Neurosci. 32, 3612–3628 (2012).

18. N. A. Steinemann, G. M. Stine, E. M. Trautmann, A. Zylberberg, D. M. Wolpert, M. N. Shadlen, Direct observation of the neural computations underlying a single decision. eLife 12 (2023).

19. Z. C. Ashwood, N. A. Roy, I. R. Stone, The International Brain Laboratory, A. E. Urai, A. K. Churchland, A. Pouget, J. W. Pillow, Mice alternate between discrete strategies during perceptual decision-making. Nat Neurosci 25, 201–212 (2022).

20. S. Pisupati, L. Chartarifsky-Lynn, A. Khanal, A. K. Churchland, Lapses in perceptual decisions reflect exploration. eLife 10, e55490 (2021).

21. J. Drugowitsch, G. C. DeAngelis, D. E. Angelaki, A. Pouget, Tuning the speed-accuracy trade-off to maximize reward rate in multisensory decision-making. eLife 4, e06678 (2015).

22. P. Simen, D. Contreras, C. Buck, P. Hu, P. Holmes, J. Cohen, Reward rate optimization in two-alternative decision making: empirical tests of theoretical predictions. J Exp Psychol Hum Percept Perform 35, 1865–1897 (2009).

23. Acquisition of decision making criteria: reward rate ultimately beats accuracy | Attention, Perception, & Psychophysics. https://link.springer.com/article/10.3758/s13414-010-0049-7.

24. N. S. Narayanan, M. Laubach, Neuronal Correlates of Post-Error Slowing in the Rat Dorsomedial Prefrontal Cortex. Journal of Neurophysiology 100, 520–525 (2008).

25. A. E. Urai, J. W. De Gee, K. Tsetsos, T. H. Donner, Choice history biases subsequent evidence accumulation. eLife 8, e46331 (2019).

26. S. Shuvaev, S. Starosta, D. Kvitsiani, A. Kepecs, A. Koulakov, “R-learning in actor-critic model offers a biologically relevant mechanism for sequential decision-making” in Advances in Neural Information Processing Systems (Curran Associates, Inc., 2020; https://papers.nips.cc/paper/2020/hash/da97f65bd113e490a5fab20c4a69f586-Abstract.html)vol. 33, pp. 18872–18882.

27. H. Dana, T.-W. Chen, A. Hu, B. C. Shields, C. Guo, L. L. Looger, D. S. Kim, K. Svoboda, Thy1-GCaMP6 Transgenic Mice for Neuronal Population Imaging In Vivo. PLOS ONE 9, e108697 (2014).

28. J. Couto, S. Musall, X. R. Sun, A. Khanal, S. Gluf, S. Saxena, I. Kinsella, T. Abe, J. P. Cunningham, L. Paninski, A. K. Churchland, Chronic, cortex-wide imaging of specific cell populations during behavior. Nat Protoc 16, 3241–3263 (2021).

29. Z. V. Guo, N. Li, D. Huber, E. Ophir, D. Gutnisky, J. T. Ting, G. Feng, K. Svoboda, Flow of Cortical Activity Underlying a Tactile Decision in Mice. Neuron 81, 179–194 (2014).

30. L. Pinto, K. Rajan, B. DePasquale, S. Y. Thiberge, D. W. Tank, C. D. Brody, Task-Dependent Changes in the Large-Scale Dynamics and Necessity of Cortical Regions. Neuron 104, 810–824.e9 (2019).

31. M. Laubach, L. M. Amarante, K. Swanson, S. R. White, What, If Anything, Is Rodent Prefrontal Cortex? eNeuro 5, ENEURO.0315-18.2018 (2018).

32. F. Barthas, A. C. Kwan, Secondary Motor Cortex: Where ‘Sensory’ Meets ‘Motor’ in the Rodent Frontal Cortex. Trends in Neurosciences 40, 181–193 (2017).

33. T.-W. Chen, N. Li, K. Daie, K. Svoboda, A Map of Anticipatory Activity in Mouse Motor Cortex. Neuron 94, 866–879.e4 (2017).

34. T. D. Hanks, C. D. Kopec, B. W. Brunton, C. A. Duan, J. C. Erlich, C. D. Brody, Distinct relationships of parietal and prefrontal cortices to evidence accumulation. Nature 520, 220– 223 (2015).

35. L. Pinto, D. W. Tank, C. D. Brody, Multiple timescales of sensory-evidence accumulation across the dorsal cortex. eLife 11, e70263 (2022).

36. M. Laubach, J. Wessberg, M. A. L. Nicolelis, Cortical ensemble activity increasingly predicts behaviour outcomes during learning of a motor task. Nature 405, 567–571 (2000).

37. C. D. Harvey, P. Coen, D. W. Tank, Choice-specific sequences in parietal cortex during a virtual-navigation decision task. Nature 484, 62–68 (2012).

38. A. J. Peters, S. X. Chen, T. Komiyama, Emergence of reproducible spatiotemporal activity during motor learning. Nature 510, 263–267 (2014).

39. J. Wang, D. Narain, E. A. Hosseini, M. Jazayeri, Flexible timing by temporal scaling of cortical responses. Nat Neurosci 21, 102–110 (2018).

40. H. Sohn, D. Narain, N. Meirhaeghe, M. Jazayeri, Bayesian Computation through Cortical Latent Dynamics. Neuron 103, 934–947.e5 (2019).

41. M. I. Leon, M. N. Shadlen, Representation of time by neurons in the posterior parietal cortex of the macaque. Neuron 38, 317–327 (2003).

42. Task-Dependent Changes in the Large-Scale Dynamics and Necessity of Cortical Regions: Neuron. https://www.cell.com/neuron/fulltext/S0896-6273(19)30731-7.

43. K. G. Thompson, K. L. Biscoe, T. R. Sato, Neuronal Basis of Covert Spatial Attention in the Frontal Eye Field. J. Neurosci. 25, 9479–9487 (2005).

44. B. B. Scott, C. M. Constantinople, A. Akrami, T. D. Hanks, C. D. Brody, D. W. Tank, Fronto-parietal Cortical Circuits Encode Accumulated Evidence with a Diversity of Timescales. Neuron 95, 385–398.e5 (2017).

45. S. A. Koay, A. S. Charles, S. Y. Thiberge, C. D. Brody, D. W. Tank, Sequential and efficient neural-population coding of complex task information. Neuron 110, 328–349.e11 (2022).

46. Z. Yang, M. Inagaki, C. R. Gerfen, L. Fontolan, H. K. Inagaki, Integrator dynamics in the cortico-basal ganglia loop underlie flexible motor timing. bioRxiv [Preprint] (2024). 10.1101/2024.06.29.601348.

47. H. K. Inagaki, S. Chen, M. C. Ridder, P. Sah, N. Li, Z. Yang, H. Hasanbegovic, Z. Gao, C. R. Gerfen, K. Svoboda, A midbrain-thalamus-cortex circuit reorganizes cortical dynamics to initiate movement. Cell 185, 1065–1081.e23 (2022).

48. J. R. de Leeuw, R. A. Gilbert, B. Luchterhandt, jsPsych: Enabling an Open-Source Collaborative Ecosystem of Behavioral Experiments. Journal of Open Source Software 8, 5351 (2023).

49. S. Rajananda, H. Lau, B. Odegaard, A Random-Dot Kinematogram for Web-Based Vision Research. Journal of Open Research Software 6 (2018).

50. S. Musall, M. T. Kaufman, A. L. Juavinett, S. Gluf, A. K. Churchland, Single-trial neural dynamics are dominated by richly varied movements. Nat Neurosci 22, 1677–1686 (2019).

51. D. H. Brainard, The Psychophysics Toolbox. Spat Vis 10, 433–436 (1997).

52. J. Peirce, J. R. Gray, S. Simpson, M. MacAskill, R. Höchenberger, H. Sogo, E. Kastman, J. K. Lindeløv, PsychoPy2: Experiments in behavior made easy. Behav Res 51, 195–203 (2019).

53. T. D. Hanks, C. Summerfield, Perceptual Decision Making in Rodents, Monkeys, and Humans. Neuron 93, 15–31 (2017).

54. R. Ratcliff, F. Tuerlinckx, Estimating parameters of the diffusion model: Approaches to dealing with contaminant reaction times and parameter variability. Psychonomic Bulletin & Review 9, 438–481 (2002).

55. M. Pachitariu, C. Stringer, M. Dipoppa, S. Schröder, L. F. Rossi, H. Dalgleish, M. Carandini, K. D. Harris, Suite2p: beyond 10,000 neurons with standard two-photon microscopy. bioRxiv [Preprint] (2017). 10.1101/061507.

